# Body size interacts with the structure of the central nervous system: A multi-center in vivo neuroimaging study

**DOI:** 10.1101/2024.04.29.591421

**Authors:** René Labounek, Monica T. Bondy, Amy L. Paulson, Sandrine Bédard, Mihael Abramovic, Eva Alonso-Ortiz, Nicole T Atcheson, Laura R. Barlow, Robert L. Barry, Markus Barth, Marco Battiston, Christian Büchel, Matthew D. Budde, Virginie Callot, Anna Combes, Benjamin De Leener, Maxime Descoteaux, Paulo Loureiro de Sousa, Marek Dostál, Julien Doyon, Adam V. Dvorak, Falk Eippert, Karla R. Epperson, Kevin S. Epperson, Patrick Freund, Jürgen Finsterbusch, Alexandru Foias, Michela Fratini, Issei Fukunaga, Claudia A. M. Gandini Wheeler-Kingshott, GianCarlo Germani, Guillaume Gilbert, Federico Giove, Francesco Grussu, Akifumi Hagiwara, Pierre-Gilles Henry, Tomáš Horák, Masaaki Hori, James M. Joers, Kouhei Kamiya, Haleh Karbasforoushan, Miloš Keřkovský, Ali Khatibi, Joo-won Kim, Nawal Kinany, Hagen Kitzler, Shannon Kolind, Yazhuo Kong, Petr Kudlička, Paul Kuntke, Nyoman D. Kurniawan, Slawomir Kusmia, Maria Marcella Laganà, Cornelia Laule, Christine S. W. Law, Tobias Leutritz, Yaou Liu, Sara Llufriu, Sean Mackey, Allan R. Martin, Eloy Martinez-Heras, Loan Mattera, Kristin P. O’Grady, Nico Papinutto, Daniel Papp, Deborah Pareto, Todd B. Parrish, Anna Pichiecchio, Ferran Prados, Àlex Rovira, Marc J. Ruitenberg, Rebecca S. Samson, Giovanni Savini, Maryam Seif, Alan C. Seifert, Alex K. Smith, Seth A. Smith, Zachary A. Smith, Elisabeth Solana, Yuichi Suzuki, George W Tackley, Alexandra Tinnermann, Jan Valošek, Dimitri Van De Ville, Marios C. Yiannakas, Kenneth A. Weber, Nikolaus Weiskopf, Richard G. Wise, Patrik O. Wyss, Junqian Xu, Julien Cohen-Adad, Christophe Lenglet, Igor Nestrašil

## Abstract

Clinical research emphasizes the implementation of rigorous and reproducible study designs that rely on between-group matching or controlling for sources of biological variation such as subject’s sex and age. However, corrections for body size (i.e. height and weight) are mostly lacking in clinical neuroimaging designs. This study investigates the importance of body size parameters in their relationship with spinal cord (SC) and brain magnetic resonance imaging (MRI) metrics. Data were derived from a cosmopolitan population of 267 healthy human adults (age 30.1±6.6 years old, 125 females). We show that body height correlated strongly or moderately with brain gray matter (GM) volume, cortical GM volume, total cerebellar volume, brainstem volume, and cross-sectional area (CSA) of cervical SC white matter (CSA-WM; 0.44≤r≤0.62). In comparison, age correlated weakly with cortical GM volume, precentral GM volume, and cortical thickness (-0.21≥r≥-0.27). Body weight correlated weakly with magnetization transfer ratio in the SC WM, dorsal columns, and lateral corticospinal tracts (-0.20≥r≥-0.23). Body weight further correlated weakly with the mean diffusivity derived from diffusion tensor imaging (DTI) in SC WM (r=-0.20) and dorsal columns (-0.21), but only in males. CSA-WM correlated strongly or moderately with brain volumes (0.39≤r≤0.64), and weakly with precentral gyrus thickness and DTI-based fractional anisotropy in SC dorsal columns and SC lateral corticospinal tracts (-0.22≥r≥-0.25). Linear mixture of sex and age explained 26±10% of data variance in brain volumetry and SC CSA. The amount of explained variance increased at 33±11% when body height was added into the mixture model. Age itself explained only 2±2% of such variance. In conclusion, body size is a significant biological variable. Along with sex and age, body size should therefore be included as a mandatory variable in the design of clinical neuroimaging studies examining SC and brain structure.

## Introduction

Knowledge about the relationship between body size (i.e., height and weight), spinal cord (SC) and brain structure is essential for a mechanistic understanding of human physiology and pathophysiology and, consequently, developing biomarkers critical for robust clinical trial designs. Besides sex and age, numerous other factors influence body size, including genetic makeup, race and ethnicity, socioeconomic and environmental factors, as well as developmental determinants. There are also diseasesaffecting physical makeup, spanning chronic conditions (i.e., anemia, asthma, celiac disease, inflammatory bowel disease, kidney or heart insufficiency), hormonal diseases (i.e., growth or thyroid hormone disbalances) and/or rare disorders such as achondroplasia and Down, Noonan or Turner syndromes.^1,2^ For example, patients diagnosed with Friedreich ataxia tend to be underweight in young age and overweight in adulthood.^3,4^ Patients with different types of mucopolysaccharidoses are known to present with a short stature.^5–7^ While neuroimaging measurements are usually compared to a healthy population, neither body height nor weight have been rigorously considered as putative confounding factors, normalization factors, and/or as variables necessary for an inter-population matching.^8–13^ Such a study design deficit can lead to bias in clinical outcomes, which applies even more explicitly to studies where the typical body size of the patients’ cohort differs from the control group. To assess the significance and importance of body size correction, we have investigated the impact of body size on structural neuroimaging measurements in the SC and brain of a healthy human population. *<u>If the</u> <u>effect is significant, future clinical research studies and trials utilizing neuroimaging should include body size as</u> <u>a potential confounding biological factor to avoid bias in clinical outcomes</u>*.

Evolutionary biology has identified links between species’ body weight, SC, and cerebral weights^14^, and between spinal canal dimensions and adjacent cord.^15,16^ Cadaveric human measurements revealed links between the cross-sectional area (CSA) of the cervical SC and cerebral weight, body height, and age.^17^ However, in vivo evidence of such a relationship between body size and central nervous system (CNS) structure is limited to a few magnetic resonance imaging (MRI) studies. In vivo CSA of the upper cervical SC (i.e., C2/3 segment) appears to be determined by both the cerebral volume and white matter (WM) content of cerebrospinal tracts.^18^ Recent exploration of the UK Biobank imaging dataset observed weak in vivo links between the CSA of the C2/3 SC segment and body height and weight, and moderate links between the CSA and brain and thalamus volumes.^19–21^ Weak correlations between body height, CSA of the SC (CSA-SC) and gray matter (GM) as well as brain volume scaling were also reported on a concurrent in vivo dataset.^22^ However, these effects disappeared when sex was controlled for.^22^ Additionally, the in vivo CSA of peripheral nerves has also been shown to moderately correlate with body height, body weight, and body mass index (BMI), but not age.^23^ Whether SC WM and GM contents are equally correlated with body size and distinct brain morphology has not been satisfactorily determined. Our **first hypothesis** was therefore that *<u>“CSA of cervical</u> <u>SC WM and GM interacts with body size and morphology of distinct brain structures”</u>*; we tested this premise by utilizing a multi-center *spine-generic* MRI dataset. The dataset allows for the separate assessment of cervical SC WM and GM morphology in a large cohort of healthy cosmopolitan volunteers with available demographic records and images of cerebral morphology.^24,25^

Myelin content is an essential characteristic of the neural tissue microstructural integrity.^26^ In the CNS, the ratio between axon diameter and diameter of the total nerve fiber (axon and myelin) is 0.6–0.7.^27^ As SC axons generally have larger diameters than axons within the brain,^28–31^ SC myelin sheaths are often also thicker, increasing the overall diameter of the myelinated axons. Thicker myelin sheaths around axons accelerate nerve conduction speed independent of the axonal diameter.^32–34^ Assuming a fairly constant axon/fiber diameter ratio,^27^ thicker myelin sheaths are therefore expected for species with larger body sizes.^33,34^ Considering intra-species variability in body size, the overall degree of SC myelination might be influenced by the body size of a given specimen. If true, the influence of body size on myelin content may be detectable in SC images sensitive to tissue microstructure, such as diffusion tensor imaging (DTI) or magnetization transfer ratio (MTR) imaging. Both DTI and MTR image contrasts are available within the *spine-generic* dataset.^24,25^ Moreover, body weight and BMI are correlated with MTR of peripheral nerves and muscles.^35^ Our **second hypothesis** was therefore that *<u>“SC microstructure, as measured using MTR and DTI, interacts with body size”</u>*.

Finally, the human brain volume and CSA-SC differ between sexes.^21,22,36,37^ It is well established that brain volume shrinks and cortical GM thickness thins with aging,^38–40^ with both processes accelerating after 45 years of age.^38,41^ However, results obtained from pathological^17,42–45^ and neuroimaging^42^ studies investigating the relationships between age and SC CSA have been less consistent. Recent high-resolution in vivo neuroimaging indeed observed weaker and slower aging effects in SC CSA than those described for brain morphology.^21,22,37^ The UK Biobank dataset already showed that physical measures, including body height and weight, strongly impact quantitative brain structural measures in a population of 40-69 years olds while adjusted for sex and age.^46^ Outside of the UK Biobank, links between body size and brain volume have been reported with inconsistent results, spanning significant relationships with a stronger height influence,^21,41,47^ or non-significant findings.^48^ Therefore, our **third hypothesis** was that: *<u>“Cerebral morphology interacts with body</u> <u>height more profoundly than with body weight and age,”</u>* We tested our hypotheses by utilizing the *spine-generic* dataset of predominantly non-elderly healthy adults and considering sex effects.

## Results

### Study cohort demography

Structural MRI data were acquired in a cohort of 267 neurologically healthy (self-reported) volunteers whose demographic data are summarized in **Table 1**. There was no significant difference in age between females and males, but body height, weight, and BMI differed (**Table 1**). All subject-specific demographic data are available at: https://github.com/spine-generic/data-multi-subject/blob/r20231212/participants.tsv. Body height and weight were significantly intercorrelated (Pearson correlation coefficient r=0.702).

**Table 1:** Demography of recruited cohort.

|  | All | Female | Male | p-value |
| --- | --- | --- | --- | --- |
| **Number of subjects** | 267 | 125 (46.82%) | 142 (53.18%) |  |
| **Age [years]** | 30.1±6.6 (19.0-56.0) | 29.4±6.4 (20.0-56.0) | 30.6±6.7 (19.0-56.0) | 0.1537 |
| **Height [cm]** | 172.1±10.0 (148.0-203.0) | 164.9±6.5 (148.0-185.0) | 178.5±8.0 (161.0-203.0) | <0.0001 |
| **Weight [kg]** | 68.3±13.4 (41.0-120.0) | 59.5±9.7 (41.0-86.0) | 76.0±11.4 (55.0-120.0) | <0.0001 |
| **BMI [kg/m<sup>2</sup>]** | 22.9±3.3 (16.6-35.5) | 21.9±3.5 (16.6-35.5) | 23.8±2.8 (18.6-35.1) | <0.0001 |

Cell values are as follows: mean ± standard deviation (minimum-maximum). P-value was derived from a two-sample t-test comparing variable distributions between females and males. BMI is the body mass index.

### Gaussianity of demographic and structural MRI data

Age demonstrated log-Gaussian distribution. Body height demonstrated neither Gaussian (p=0.0089), nor log-Gaussian distribution (p=0.0259). Body weight, BMI, all CSA measurements, all SC DTI measurements and all brain morphological measurements demonstrated Gaussian distributions. All SC MTR measurements demonstrated neither Gaussian (p<0.0086) nor log-Gaussian (p<0.0009) distributions.

### Body size interacts with the structure of spinal cord white matter

The following CSA measurements were averaged from cervical C3-4 segments (see Methods for details). CSA of SC (CSA-SC) was moderately correlated with body height (r=0.355, **Fig. 1**, **Table 2**), and this correlation strength was higher for the CSA-WM subregion (r=0.437, **Fig. 1**, **Table 2**). CSA-SC and CSA-WM demonstrated minimal differences between scanner manufacturers (**Fig. 1**). Thus, the same correlation patterns for height were preserved even when manufacturer-specific averages of CSA-SC or CSA-WM were subtracted from corresponding CSA measurements prior to the correlation analysis in order to normalize data across scanners (**Table 2**). The correlation between body height and CSA-SC/CSA-WM remained significant even when the dataset was split into males and females (**Table 2**). Body weight was more weakly correlated with CSA-SC (r=0.261) and CSA-WM (r=0.274). In addition, this correlation was not significant when the dataset was split into males and females (**Supplementary Table 1**). CSA-GM was not correlated with body size (**Fig. 1**, **Supplementary Table 1**). The CSA-GM measured on Philips scanners demonstrated a lower mean offset than for data obtained on Siemens and GE scanners (**Fig. 1**; p<0.0001). Neither CSA measurement (i.e, SC, WM, GM) was correlated with age (**Fig. 1**, **Supplementary Table 1**). Overall, body height is the demographic variable driving the impact on CSA-WM and explaining the majority of demography-related variability in CSA measurements **(Fig. 2b**, **Table 3**).

**Figure 1:**
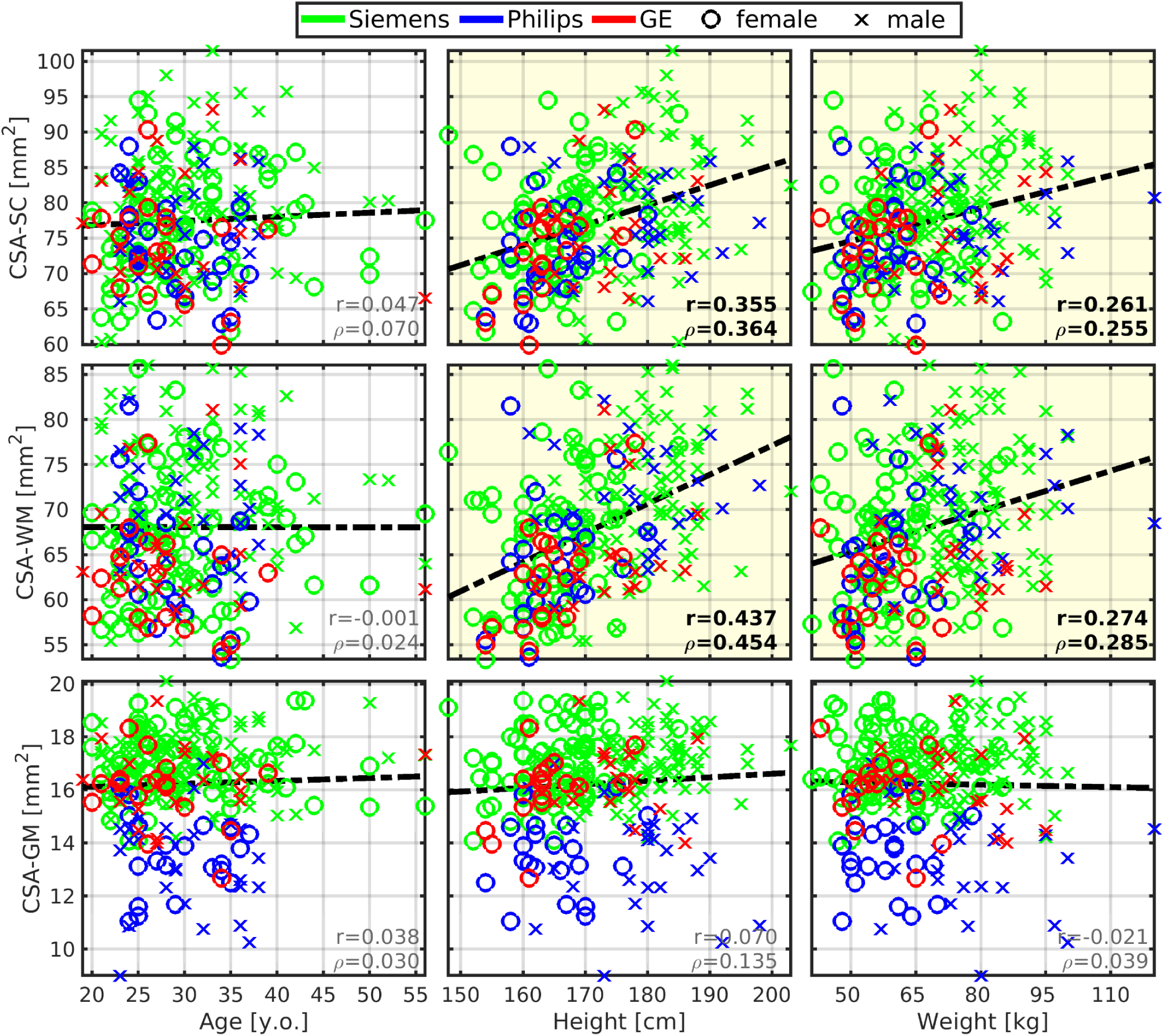
Cross-sectional area of spinal cord white matter correlates with body height and weight. *<u>Abbreviations:</u>* CSA - cross-sectional area; SC - spinal cord; WM - white matter; GM - gray matter; r - Pearson correlation coefficient; ⍴ - Spearman correlation coefficient. All spinal cord measurements were averaged from cervical C3-4 levels. Regression lines (i.e., the dashed black lines) were estimated from all available data points. Plots with statistically significant correlation (p_FWE_<0.05) are highlighted with yellow background, and corresponding r and ⍴ values are highlighted with black bold font.

**Figure 2:**
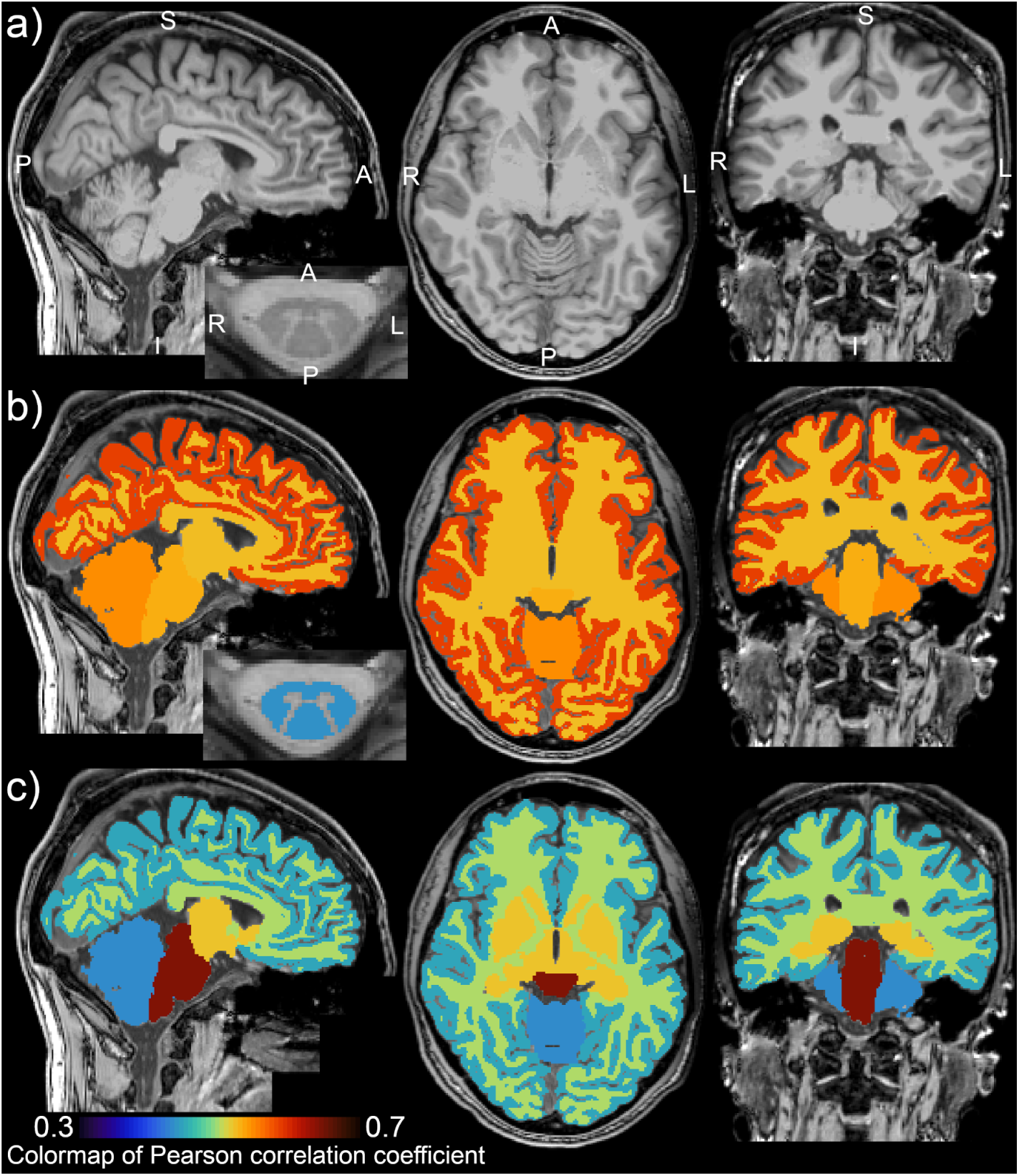
Interactions between body height and morphology of the central nervous system. **Panel a)** Representative image of brain and spinal cord (SC) anatomy. The brain scan shows cortical gray matter (GM), cerebral white matter (WM), subcortical GM structures, brainstem and cerebellum. The axial SC scan shows the WM and GM anatomy at the C3/C4 level. Image orientation is described in **panel a)**: A - anterior, P - posterior, S - superior, I - Inferior, L - left and R - right. **Panel b)** Pearson correlation coefficient between body height and (i) cortical GM volume; (ii) cerebral WM volume; (iii) subcortical GM structure volume; (iv) brainstem volume; (v) cerebellar volume; and (vi) cross-sectional area (CSA) of cervical SC WM at C3/C4 level. The colormap for the correlation values is shown in the left bottom corner of the figure. All correlations are significant (p_FWE_<0.05). Regarding the investigated list of structures, body height demonstrated the strongest correlation with the cortical GM volume. **Panel c)** Pearson correlation coefficient between the CSA of cervical WM at C3/C4 level and (i) cortical GM volume; (ii) cerebral WM volume; (iii) subcortical GM structure volume; (iv) brainstem volume; and (v) cerebellar volume. The colormap for the correlation values is shown in the left bottom corner of the figure. All correlations are significant (p_FWE_<0.05). The correlation map shows a descending gradient from the brainstem through subcortical GM structures and cerebral WM to cortical GM. The gradient may be driven by the increasing distance to the cervical SC level and decreasing relative volume of common tract pathways. The cerebellum shows the lowest, yet significant, correlation level. This finding may be explained by the fact that the cerebrum is more strongly and directly interconnected to the peripheral nervous system via SC than the cerebellum, with spinocerebellar tracts as the primary direct connections.^50,51^

**Table 2:** Pearson correlation coefficients between body size, age, spinal cord structure, and brain structure, and post-hoc sex-effects in the correlation analysis. *<u>Abbreviations:</u>* CSA - cross-sectional area; SC - spinal cord; WM - white matter; GM - gray matter; Vol - volume. The correlation analysis on non-normalized data identified a list of variable pairs with a correlation coefficient of p_FWE_<0.05. The final list here only selects the variable pairs with the significant post-hoc Pearson correlation coefficient (uncorrected p<0.05) in at least one sex-specific sub-dataset (i.e., female and/or male). Insignificant correlation coefficients, that did not meet the post-hoc condition uncorrected p<0.05, are written with gray font. Positive correlation coefficients (p<0.05) are visualized as a yellow-orange-pink-red color shade of the table background. Negative correlation coefficients (p<0.05) are visualized as a light blue-blue color shade of the table background. CSA was measured as averages between C3-C4 segments. DTI and MTR were calculated as averages between C2-C5 segments. The column denoted *“Original values”* reports correlation coefficients for raw measurements with no normalization procedure prior to the correlation analysis. The column denoted *“Manufacturer-specific normalized SC measures”* reports correlation coefficients for SC structural measurements, which were normalized to zero mean for each scanner manufacturer before correlation analysis. Empty cells in the right half of the table represent combinations where no updated correlation coefficients were measured, because the utilized normalization of SC structural measurements had no effect on these correlation coefficients. Brain structural measurements were not considered necessary to normalize as we did not observe strong scanner-related effects in brain macrostructural measurements.

| Grouping Variable | Correlated variable | Original values (no data normalization) |  |  | Manufacturer-specific normalized SC measures |  |  |
| --- | --- | --- | --- | --- | --- | --- | --- |
|  |  | All | Female | Male | All | Female | Male |
| HEIGHT | BrainGMVol | 0.622 | 0.321 | 0.446 |  |  |  |
|  | BrainVol | 0.612 | 0.274 | 0.408 |  |  |  |
|  | CorticalGMVol | 0.583 | 0.252 | 0.448 |  |  |  |
|  | CerebellumVol | 0.546 | 0.411 | 0.218 |  |  |  |
|  | BrainStemVol | 0.531 | 0.310 | 0.258 |  |  |  |
|  | CorticalWMVol | 0.523 | 0.157 | 0.312 |  |  |  |
|  | SubCortGMVol | 0.522 | 0.107 | 0.287 |  |  |  |
|  | PrecentralGMVol | 0.495 | 0.092 | 0.418 |  |  |  |
|  | CSA-WM | 0.437 | 0.295 | 0.303 | 0.422 | 0.285 | 0.268 |
|  | PostcentralGMVol | 0.434 | 0.121 | 0.369 |  |  |  |
|  | CSA-SC | 0.355 | 0.323 | 0.230 | 0.344 | 0.319 | 0.205 |
| WEIGHT | BrainStemVol | 0.431 | 0.119 | 0.191 |  |  |  |
|  | MD-SC-WM | -0.200 | -0.022 | -0.191 | -0.252 | -0.108 | -0.206 |
|  | MTR-SC-WM | -0.225 | -0.374 | -0.221 | -0.221 | -0.331 | -0.217 |
| AGE | CorticalGMVol | -0.213 | -0.357 | -0.258 |  |  |  |
|  | PrecentralGMVol | -0.205 | -0.326 | -0.232 |  |  |  |
|  | Cortical Thickness | -0.274 | -0.278 | -0.277 |  |  |  |
| CSA-WM | BrainStemVol | 0.640 | 0.624 | 0.547 | 0.590 | 0.580 | 0.467 |
|  | BrainVol | 0.524 | 0.392 | 0.459 | 0.507 | 0.379 | 0.426 |
|  | SubCortGMVol | 0.511 | 0.257 | 0.524 | 0.488 | 0.248 | 0.474 |
|  | CorticalWMVol | 0.503 | 0.315 | 0.471 | 0.501 | 0.339 | 0.447 |
|  | BrainGMVol | 0.481 | 0.341 | 0.406 | 0.445 | 0.286 | 0.359 |
|  | CorticalGMVol | 0.449 | 0.283 | 0.390 | 0.412 | 0.227 | 0.346 |
|  | CerebellumVol | 0.433 | 0.501 | 0.179 | 0.441 | 0.508 | 0.186 |
|  | ThalamusVol | 0.431 | 0.244 | 0.398 | 0.451 | 0.279 | 0.408 |
|  | PrecentralGMVol | 0.420 | 0.235 | 0.382 | 0.370 | 0.178 | 0.317 |
|  | PostcentralGMVol | 0.389 | 0.243 | 0.356 | 0.345 | 0.181 | 0.311 |
|  | PrecentralG Thickness | 0.252 | 0.249 | 0.209 | 0.146 | 0.142 | 0.087 |
| CSA-SC | BrainStemVol | 0.575 | 0.622 | 0.492 | 0.519 | 0.573 | 0.409 |
|  | BrainVol | 0.419 | 0.337 | 0.368 | 0.393 | 0.311 | 0.327 |
|  | SubCortGMVol | 0.417 | 0.207 | 0.449 | 0.388 | 0.198 | 0.392 |
|  | CorticalWMVol | 0.432 | 0.291 | 0.422 | 0.411 | 0.280 | 0.384 |
|  | BrainGMVol | 0.357 | 0.265 | 0.287 | 0.324 | 0.225 | 0.242 |
|  | CorticalGMVol | 0.320 | 0.195 | 0.262 | 0.290 | 0.159 | 0.225 |
|  | CerebellumVol | 0.356 | 0.508 | 0.109 | 0.342 | 0.493 | 0.086 |
|  | ThalamusVol | 0.335 | 0.176 | 0.322 | 0.338 | 0.201 | 0.308 |
|  | PrecentralGMVol | 0.307 | 0.159 | 0.273 | 0.275 | 0.132 | 0.228 |
|  | PostcentralGMVol | 0.240 | 0.146 | 0.184 | 0.218 | 0.120 | 0.160 |
|  | PrecentralG Thickness | 0.211 | 0.180 | 0.201 | 0.144 | 0.125 | 0.117 |
| CSA-GM | BrainStemVol | 0.390 | 0.444 | 0.480 | 0.335 | 0.440 | 0.375 |
|  | BrainVol | 0.221 | 0.250 | 0.298 | 0.186 | 0.219 | 0.254 |
|  | SubCortGMVol | 0.217 | 0.093 | 0.392 | 0.197 | 0.119 | 0.343 |
|  | CorticalWMVol | 0.268 | 0.275 | 0.355 | 0.217 | 0.215 | 0.301 |

**Table 3:**
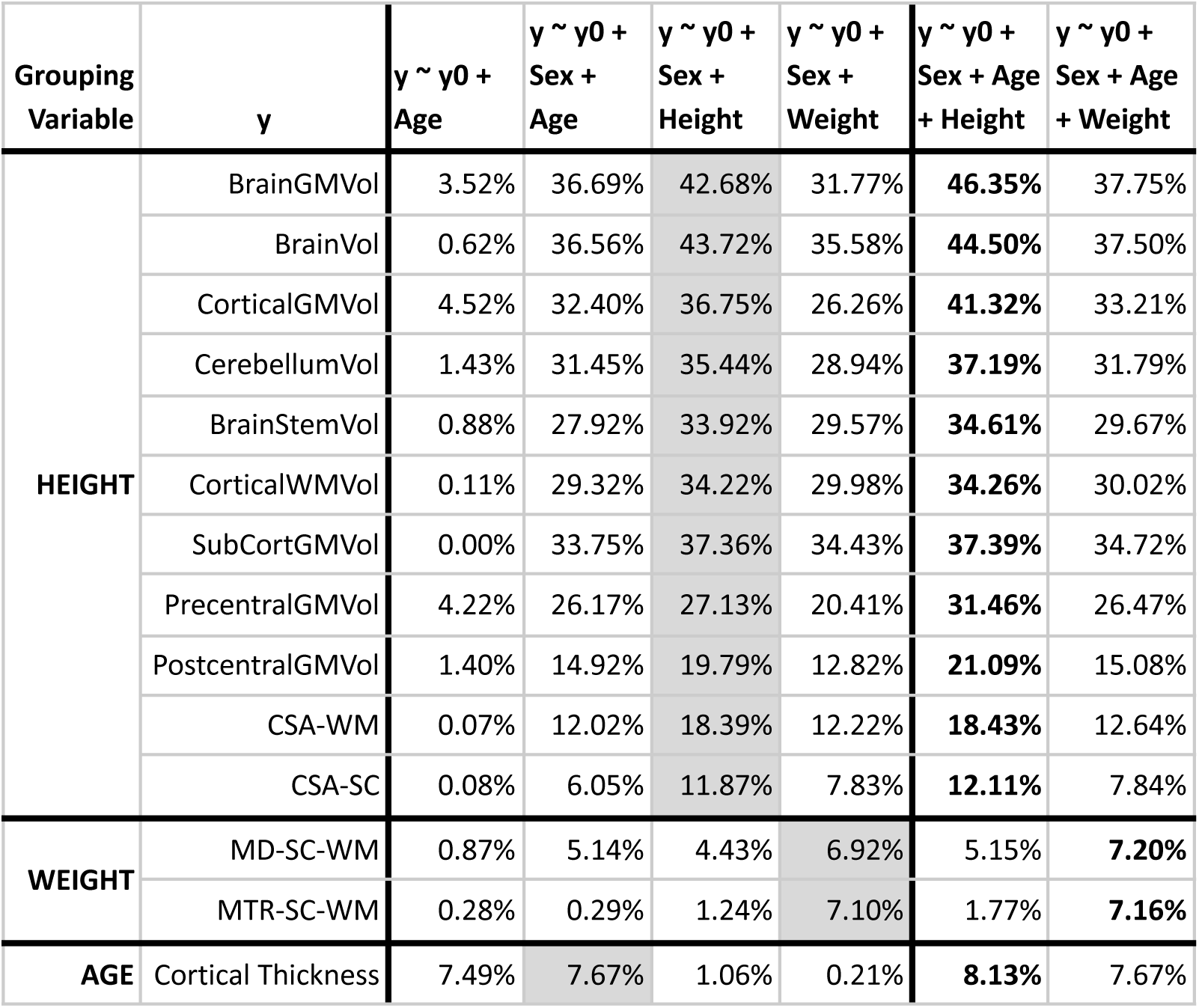
Coefficients of determination (R^2^) of regression models explaining a variable y as a linear mixture of age, body height and body weight. Gray background highlighted R^2^ coefficients are those whose single demographic variable (age, height or weight) explained the most of the neuroimaging data variance when accounting for sex effects. Except for cortical thickness, body size explains a significantly higher portion of variance in brain and spinal cord structural measurements than age. Bold highlighted R^2^ coefficients identifies models which explained most of the neuroimaging data variance by given demographic variables. *<u>Abbreviations:</u>* CSA - cross-sectional area; SC - spinal cord; WM - white matter; GM - gray matter; Vol - volume.

DTI- and MTR-derived microstructural measurements were averaged from cervical C2-5 levels (see Methods for details). GE-scanner-derived DTI and MTR measurements significantly differed from Siemens and Philips scanners (**Fig. 3**, p<0.0001). Therefore, GE scanner microstructural measurements (13.87% of the dataset) were not included in correlation analyses that did not use manufacturer-specific normalized microstructural values (**Table 2**). Body weight was weakly correlated with mean diffusivity (MD) in the WM region (r=-0.200, **Fig. 3**, **Table 2**, **Fig. S1**) and bilateral dorsal columns (DC, r=-0.207, **Fig. S2**). Body weight was not significantly correlated to MD for females (**Table 2**). No investigated DTI measures (i.e., MD, fractional anisotropy - FA or radial diffusivity) were correlated to body size when extracted from the GM region (**Fig. S3**) or bilateral lateral corticospinal tracts (LCST; **Fig. S4**). CSA-WM and SC FA were weakly correlated in DC (r=-0.247) and LCST (r=-0.224). Body weight was weakly correlated to MTR in WM (r=-0.225, **Fig. 3**) DC (r=-0.231, **Fig. S5**) and LCST (r=-0.200, **Fig. S5**), and not correlated to MTR in GM (**Fig. S5**). The correlation between body weight and MTR remained significant, even when the dataset was split into males and females (**Table 2**). When the dataset was normalized for each manufacturer and values from GE scanners were included in the analysis, the correlation values remained almost identical (**Table 2**). This finding signifies that the observed effect remained identical but had slightly higher power due to the larger sample size (added 37 samples; +13.87%). The correlation analysis revealed no aging effects in DTI (-0.004≥r≥-0.099) or MTR (-0.047≥r≥-0.094) measures (**Fig. S1-5**) in our sample. However, the exploratory principal component analysis showed small effects in mutual covariance (**Fig. 7d**). Linear regression analysis showed that body weight explained the majority of the demography-related variance in our young adult sample DTI and MTR measurements (**Table 3**).

**Figure 3:**
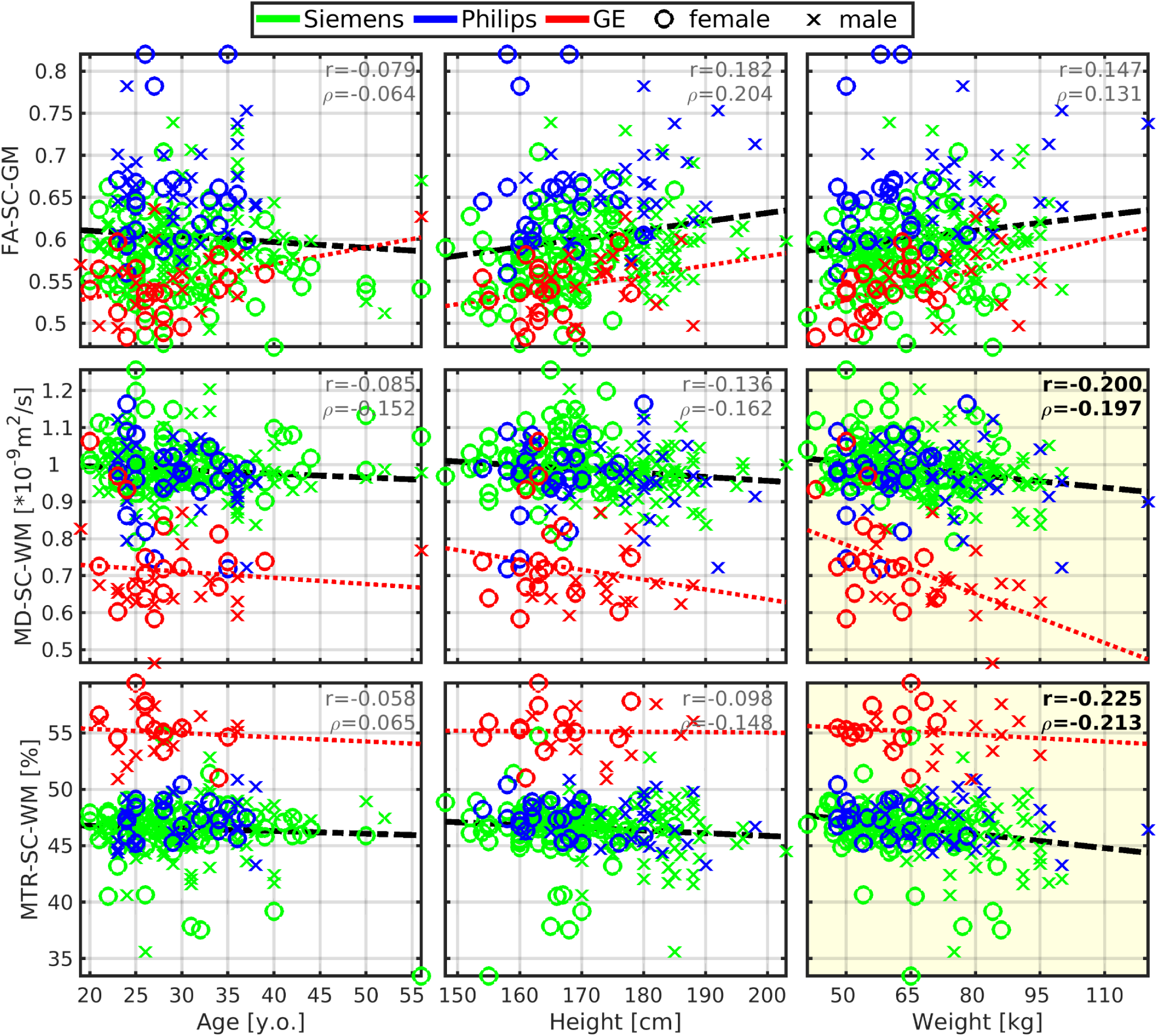
Mean diffusivity and magnetization transfer ratio in spinal cord white matter correlates with body weight. *<u>Abbreviations:</u>* GM - gray matter; WM - white matter; SC - spinal cord; FA - fractional anisotropy; MD - mean diffusivity; MTR - magnetization transfer ratio; r - Pearson correlation coefficient; ⍴ - Spearman correlation coefficient. All spinal cord measurements were averaged from cervical C2-5 levels. Black dashed regression lines were estimated from the Siemens and Philips scanners’ data points. Red dotted regression lines were estimated from the GE scanner’s data points. Plots with statistically significant correlation (p_FWE_<0.05) are highlighted with yellow background, and corresponding r and ⍴ values are highlighted with black bold font.

### Body height and age interact with brain morphology

Body height was moderately correlated with several cerebral volumes (r=0.54±0.06; 0.434≤r≤0.622), i.e., volumes of the brain, brain GM, cortical GM, cortical WM, subcortical GM, thalamus, cerebellum, brainstem, precentral GM and postcentral GM (**Fig. 4**, **Fig. 5a**). The vast majority of correlations with body height remained significant even after the dataset split to males and females, except for the volumes of cortical WM, subcortical GM, precentral GM and postcentral GM in females (**Table 2**). The body height interacted most profoundly with the cortical GM volume (**Fig. 2b**).

**Figure 4:**
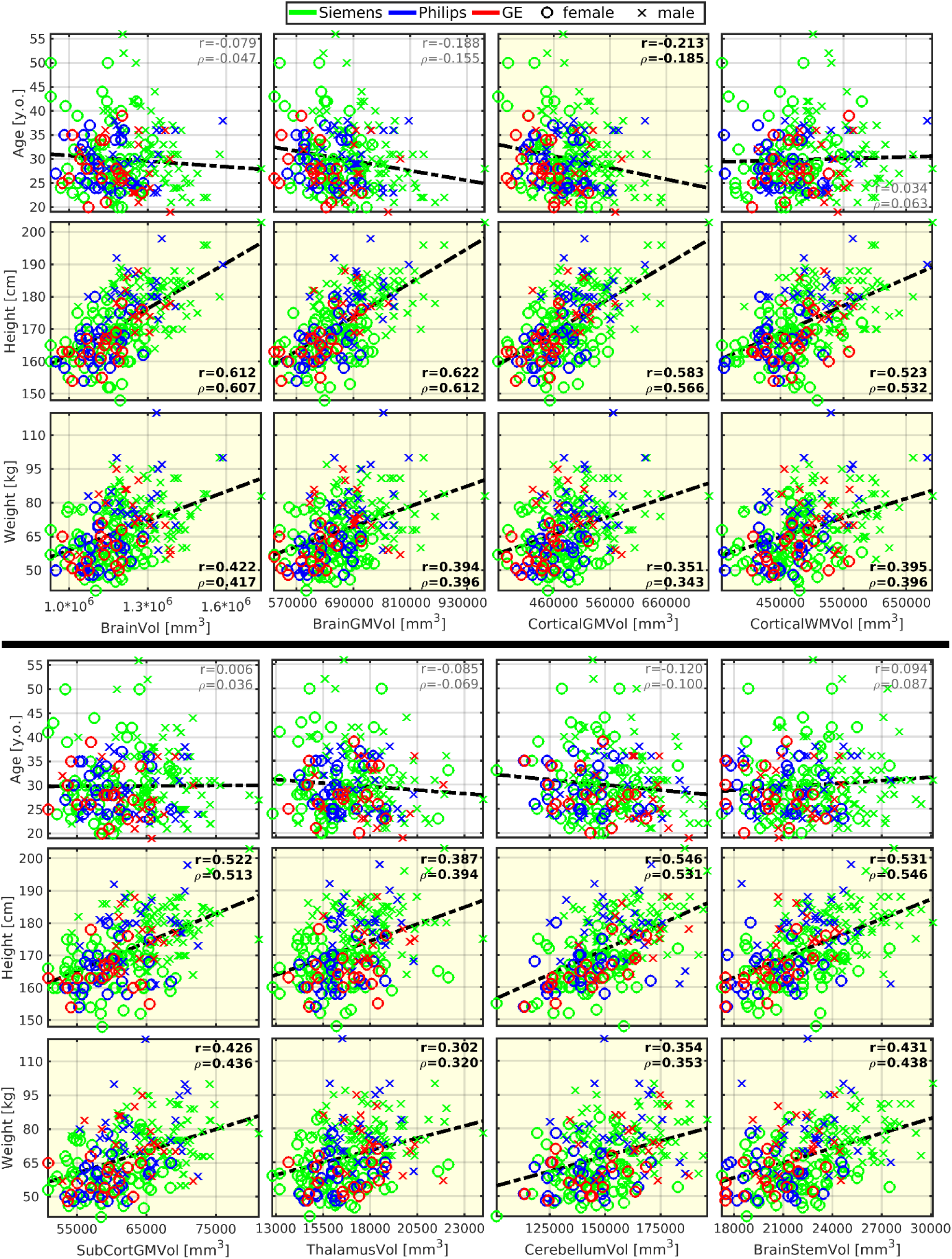
Brain morphology strongly correlates with body size but weakly with age. *<u>Abbreviations:</u>* GM - gray matter; WM - white matter; Vol - volume; SubCort - subcortical; r - Pearson correlation coefficient; ⍴ - Spearman correlation coefficient. Regression lines (i.e., the dashed black lines) were estimated from all available data points. Plots with statistically significant correlation (p_FWE_<0.05) are highlighted with yellow background, and corresponding r and ⍴ values are highlighted with black bold font.

**Figure 5:**
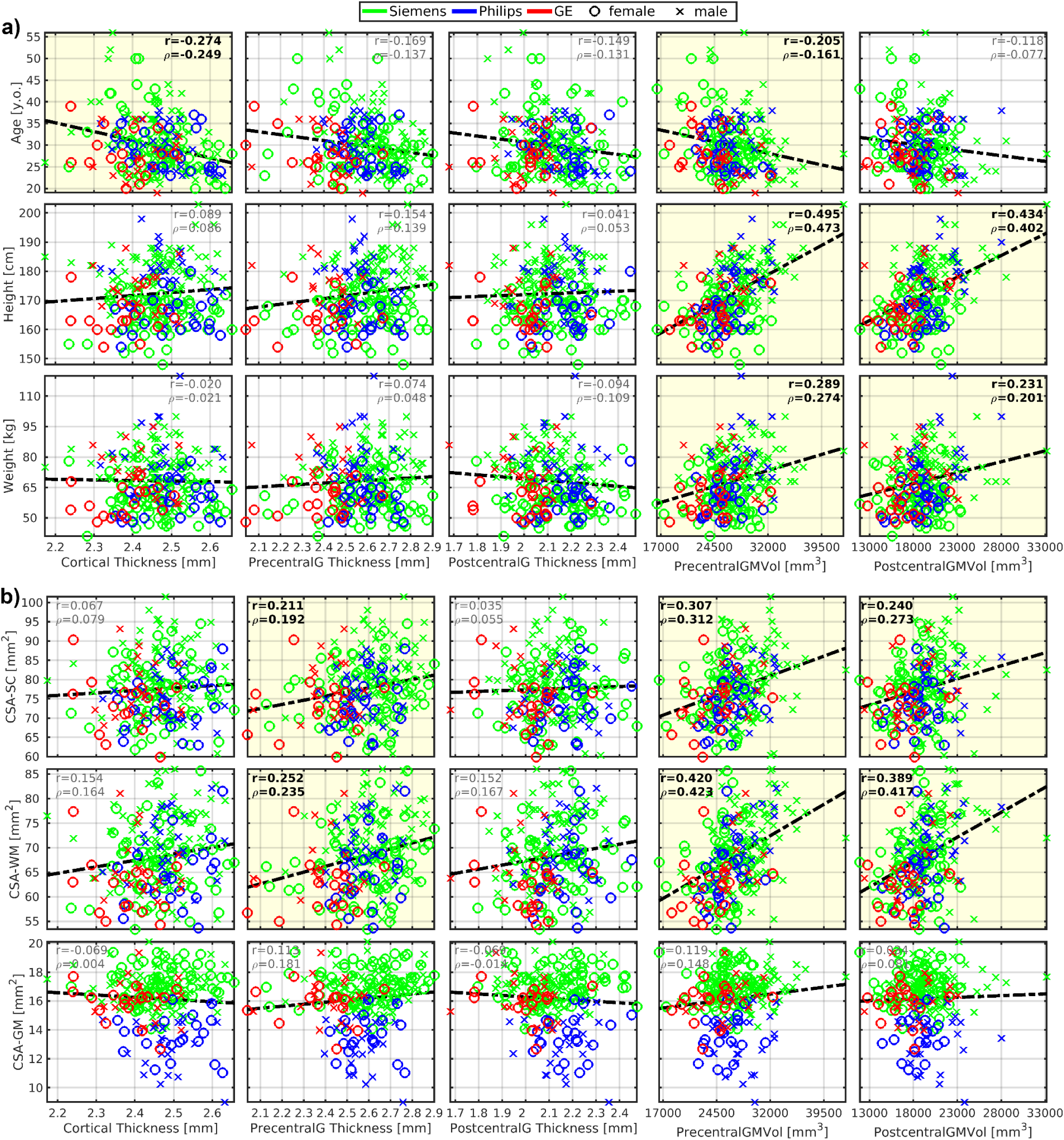
Cortical morphology correlates with body size, age, and cross-sectional area of the spinal cord white matter. *<u>Abbreviations:</u>* CSA - cross-sectional area; SC - spinal cord; WM - white matter; GM - gray matter; PrecentralG - precentral gyrus; PostcentralG - postcentral gyrus; Vol - volume; r - Pearson correlation coefficient; ⍴ - Spearman correlation coefficient. Regression lines (i.e., the dashed black lines) were estimated from all available data points. Plots with statistically significant correlation (p_FWE_<0.05) are highlighted with yellow background, and corresponding r and ⍴ values are highlighted with black bold font. **a)** Graphs demonstrate correlations with body size and age. **b)** Graphs demonstrate correlation with CSA measured in the SC region as averages from cervical C3-4 levels.

Body weight demonstrated weaker correlations in all the cortical regions that moderately correlated with the body height (r=0.37±0.07, **Fig. 4**, **Fig. 5a**, **Supplementary Table 1**). But, the only significant correlation, which survived the dataset split to males and females, was with brainstem volume in males (**Table 2**).

Body height or body weight were not correlated with total cortical, precentral gyrus, and postcentral gyrus thicknesses (**Fig. 5a**).

As expected, a weak manifestation of age-related cortical GM atrophy was observed in volume (r≈-0.213) and thickness (r=-0.274) measures (**Fig. 4**, **Fig. 5a**). The aging GM atrophy effects remained significant after the dataset split to males and females (**Table 2**).

Most importantly, the magnitude of linear dependence between brain morphology and body height was about 2- to 3-fold compared to the effects of age (**Table 2**, **Fig. 4**, **Fig. 5a**). Moreover, the young adult dataset showed that body height explains more, pathology unrelated, variance in brain volumetry than age and sex (**Table 3**). Contrary, cortical thickness variance was associated predominantly with age (**Table 3**).

### Cross-sectional area of spinal cord white matter interacts with brain morphology

CSA-SC (r=0.38±0.09; 0.240≤r≤0.575) and CSA-WM (r=0.48±0.07; 0.389≤r≤0.640) were moderately correlated with the investigated brain volumes, i.e., volumes of the brain, brain GM, cortical GM, cortical WM, subcortical GM, thalamus, cerebellum, brainstem, precentral GM and postcentral GM (**Fig. 5b**, **Fig. 6**, **Table 2**). Compared to CSA-SC, correlation strengths were higher for CSA-WM (**Fig. 5b**, **Fig. 6**, **Table 2**). CSA-GM was weakly correlated with the volume of the brain, cortical WM, subcortical GM, and brainstem, but the strength of these correlations was half weaker than those observed for CSA-SC and CSA-WM (**Fig. 6**, **Table 2**). All CSA-WM and most of the other observed correlations remained significant after the dataset split to females and males (**Table 2**) or when SC data were normalized (zero mean) for each manufacturer prior to correlation analysis (**Table 2**). CSA-WM was the primary marker defining the correlations with the brain volumes. There was a descending gradient of the CSA-WM correlation from the brainstem to subcortical GM and then cortical WM to the cortical GM (**Fig. 2c**). All these correlations were higher than the correlation with the volume of the cerebellum (**Fig. 2c**). Yet, even the correlation between CSA-WM and cerebellum volume was significant **(Fig. 2c**, **Fig. 6**).

**Figure 6:**
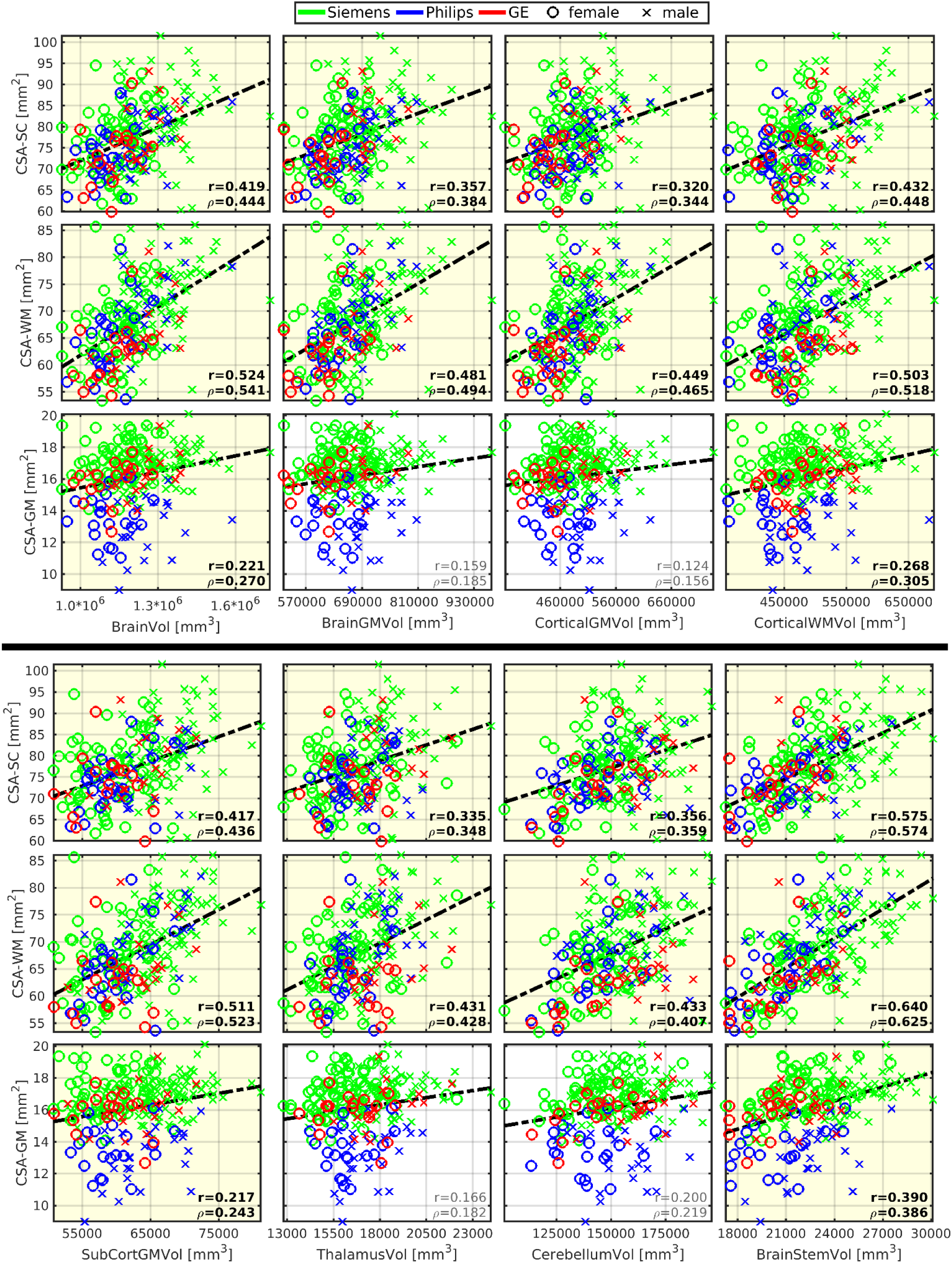
Brain morphology correlates with spinal cord morphology. *<u>Abbreviations:</u>* CSA - cross-sectional area; SC - spinal cord; WM - white matter; GM - gray matter; Vol - volume; SubCort - subcortical; r - Pearson correlation coefficient; ⍴ - Spearman correlation coefficient. All SC measurements were averaged from cervical C3-4 levels. Regression lines (i.e., the dashed black lines) were estimated from all available data points. Plots with statistically significant correlation (p_FWE_<0.05) are highlighted with yellow background, and corresponding r and ⍴ values are highlighted with black bold font.

CSA-SC (r=0.211) and CSA-WM (r=0.252) were weakly correlated with the thickness of the precentral gyrus (**Fig. 5b**). The correlations remained significant after the dataset split to females and males (**Table 2**). However, the correlations disappeared when SC data were normalized before correlation analysis (**Table 2**). CSA-GM was not correlated with any utilized cortical thickness measurement (**Fig. 5b**).

### Brain morphology and spinal cord microstructure are not related

No correlations were detected between SC WM/GM microstructure and cerebral volumes (i.e., total brain, brain GM, cortical GM, cortical WM, subcortical GM, thalamus, cerebellum, brainstem, precentral GM and postcentral GM) or cortical thickness (**Fig. S6-7**), and between thickness measurements and DTI/MTR measurements, even if the SC ROIs were limited to the bilateral LCST or DC (**Fig. S8-10**).

### Scanner-related effects on SC structural measurements

CSA-SC and CSA-WM offsets differed minimally between manufacturers (**Fig. 1**, **Fig. 5b**, **Fig. 6**). CSA-GM measurements on Philips scanners were significantly lower than CSA-GM measurements from Siemens and GE scanners (**Fig. 1**, **Fig. 5b**, **Fig. 6**, p<0.0001). Additional discussion about this specific CSA-GM issue can be found in Cohen-Adad et al. (2021)^25^. Data normalization before correlation analysis mainly decreased the correlation strengths in all CSA measures (without normalization: r=0.348±0.127; with normalization r=0.313±0.128; **Table 2**; paired t-test p<0.0001). This finding underlines the importance of adjusting for scanner-related variability in CSA measurements to minimize risks of false positive results due to scanner-related data trends.

All microstructural measurements obtained with GE scanners showed significant offsets compared to those from Siemens and Philips scanners (**Fig. 3**, **Fig. S1-10**, p<0.0001). The differences had a direct impact on correlation analyses. Therefore, we performed correlation analyses of original values without GE values and correlation analyses of normalized values utilizing all scanners’ data. Correlation analyses were stable and comparable in the magnitude of correlation coefficients for MTR (**Table 2**) and MD (**Table 2**). The normalized correlation analysis provided higher statistical power due to the larger sample size. Additionally, if we utilized GE data (**Fig. 3**) in the correlation analysis without normalization, the resulting correlation coefficients for MD-SC-WM and MTR-SC-WM in **Table 2** would be substantially lower.

### Minimal impact of degenerative cervical spinal cord compression on correlation analysis

We excluded one participant with severe degenerative cervical SC compression, providing outliers in SC structural measurements. However, the *spine-generic* database identifies an additional 61 subjects with mild degenerative compression and 2 subjects with severe degenerative compression and radiological myelopathy.^49^ Analysis power decreased, but minimal nuances were detected in correlation coefficients when tested separately on subjects without or with degenerative cervical SC compression (see **Supplementary Slides**). Thus, we conclude that SC compression had a minimal impact on the current study outcomes.

### Principal component analysis (PCA) reveals body-SC-brain structural links

We subtracted manufacturer-specific average values from all SC structural measurements prior to the exploratory analysis via PCA. PCA did not include DTI and MTR measurements from bilateral LCST and DC, as the WM region provided analogic observations. Cerebral volumes, CSA-WM, and body height formed the first principal component (PC1), characterizing 43.65% of data variance (**Fig. 7a**). CSA-SC and body weight were close, yet separated from the PC1 cluster (**Fig. 7a**). This finding supports the previously observed body height and CSA-WM dominance in the observed effects (**Fig. 1**, **Fig. 2**, **Fig. 4**, **Fig. 5**, **Fig. 6**, **Table 2**). Cortical thickness, MD-SC-WM, and age (negative effect) formed the second principal component (PC2), characterizing 12.32% of data variance (**Fig. 7a**) and presenting predominantly negative aging effects in the thickness measures. The PC1-PC3 projection showed that the PC3 characterizes about 8.91% of data variance, predominantly explained by CSA-GM, MD-SC-WM, and FA-SC-GM (negative effect), i.e., a link between SC GM morphology and SC microstructure (**Fig. 7b**). The PC2-PC3 projection verified that the cortical thickness variability predominantly forms PC2. In contrast, the PC3 is predominantly formed by SC DTI and CSA-GM (**Fig. S11a**). The PC4 explained 5.62% of unique SC DTI microstructural data variance, which is not present in other investigated modalities and investigated demographic measures (**Fig. 7c**). PC5 showed positive effects of body weight and age on MD-SC-WM, and negative effects of body weight and age on MTR-SC-WM, FA-SC-GM, and CSA-GM. These effects explained about 4.77% of data variance (**Fig. 3**, **Fig. 7d**). Simultaneously, the PC projections suggest that the impact of body weight on MTR- and DTI-derived microstructure metrics might be ≈5% (**Fig. 3**, **Fig. 7**). That quite follows the result of 7% explained variance in regression analysis (**Table 3**). However, the positive effects of body weight on MD-SC-WM contradicts our observation of weak negative correlation between body weight and SC MD (**Fig. 3**). PC3 and PC5 showed clear evidence that CSA-GM morphology and SC microstructure are linked, yet unrelated to cerebral and SC WM morphology (**Fig. 7b,d**, **Fig. S11a**). In summary, PCA analysis explained 75.27% of data variance, meaning that nearly 25% is unexplained (**Fig. S11b**).

**Figure 7:**
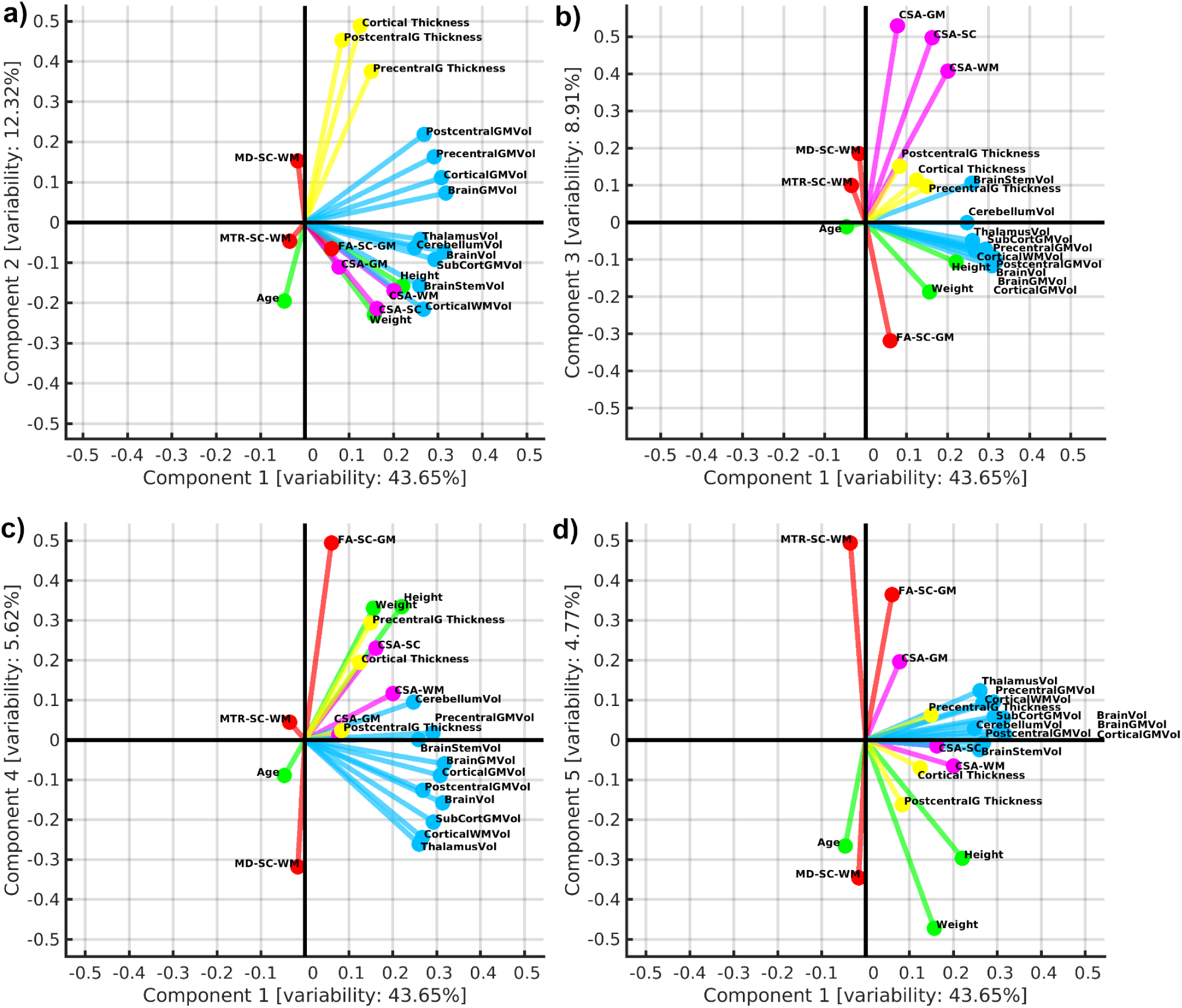
Exploratory visualization using biplot projections of principal components. **a)** biplot projection of 1^st^ and 2^nd^ principal components (PCs); **b)** biplot projection of 1^st^ and 3^rd^ PCs; **c)** biplot projection of 1^st^ and 4^th^ PCs; **d)** biplot projection of 1^st^ and 5^th^ PCs. Variable vectors are visualized in each biplot projection with a color-coding characteristic for a corresponding variable group. Variable name abbreviations and variable color codings are described as follows. *<u>Variable abbreviations:</u>* MD - mean diffusivity; RD - radial diffusivity; MTR - magnetization transfer ratio; SC - spinal cord; WM - white matter; GM - gray matter; CSA - cross-sectional area; Vol - volume; PrecentralG - precentral gyrus; PostcentralG - postcentral gyrus. *<u>Variable color coding:</u>* demography - green; cerebral volumes - light blue; cortical thicknesses - yellow; SC morphometry - magenta; SC WM microstructure - red. *<u>How to read a biplot:</u>*The overall domain of each component axis is <-1,1>. Each variable is characterized as a vector of magnitude in the range of <0,1> in the biplot space. Angle 0° between the component axis and variable vector with magnitude 1 (or between two variable vectors both with magnitude 1) is proportional to Pearson correlation coefficient 1. Under the same vector magnitude circumstances, an angle of 180° equals Pearson correlation coefficient -1, and angles of 90° and 270° equal Pearson correlation coefficient 0. The lower magnitude of variable vectors proportionally decreases the overall linear dependence between vector angles close to 0° or 180°, respectively. Similarly, angle deviation from 0° or 180° also decreases the level of linear dependence between pairs of vectors in the biplot.

## Discussion

The current study, using the *spine-generic* dataset, presents unique multi-center in vivo evidence about adult human cervical SC and brain, and emphasizes the following findings:

***(i)*** Body height correlates moderately with SC WM and brain morphology, improves explanation of demography-related variance in such structural measurements from 26±10% (range 6-37%) to 33±11% (range 12-46%) in young adults, and underlines the impact of such pathology unrelated variability in structural neuroimaging data.
***(ii)*** The expected aging effects^21,22,38–41^ explain minimal amounts of SC and brain structural data variance (2±2%) in young adults except cortical thickness (8%).
***(iii)*** Body height predominantly impacts the cortical GM volume (**Fig. 2b**) and may even define the overall brain GM volume.
***(iv)*** Body weight correlates weakly with SC WM MTR, which is influenced by myelin content.
***(v)*** Body weight correlates weakly with SC WM microstructure assessed with DTI MD.
***(vi)*** Body weight explains ≈5-7% of DTI and MTR data variance.
***(vii)*** SC WM DTI and MTR explain a significant portion of examined dataset variance (≈14-19%) and are nearly orthogonal to most macrostructural measurements, except for the CSA-GM.
***(viii)*** Subcortical and cortical GM volumes are correlated with CSA-WM more profoundly than the cerebellar volume with a descending correlation gradient from the brainstem toward cortical GM (**Fig. 2c**).
***(ix)*** Cortical WM, subcortical GM, and brainstem volumes correlate with CSA-GM but much less profoundly than CSA-WM.
***(x)*** Cortical thickness of the precentral cortex correlates weakly with CSA-WM.
***(xi)*** We highlight the importance of considering the scanner-related effects present in SC imaging data.^24,25^
***(xii)*** We confirm significant relationships between body size, brain volumes/weight, and CSA-SC in line with previously reported results.^17,21,47^

### Practical impact of the current study in clinical neuroimaging study designs

MRI of SC structure is emerging in clinical research of neurodegenerative diseases and SC injuries.^52–63^ Microstructural SC MRI of neural tissue integrity aims to understand pathophysiological changes at the subclinical or presymptomatic stage.^9,64–66^ Quantitative MRI has made significant advances over the past two decades for brain imaging,^67–86^ but is still in its early development stage when it comes to SC imaging. Sex- and age-matching are critical for any clinical neuroimaging study. Yet, we are proposing that mismatched variability in body size may influence imaging outcomes more profoundly than mismatched variability in age. Persistent marginal impact of body stature on brain structural and functional neuroimaging outcomes in the early elderly population^46,87^ further underlines the importance of our proposal. Therefore, body size needs to be considered in the rigor of future neuroimaging studies focusing on between-group differences in brain or SC structure to secure and guarantee the reproducibility of results. It has not been a common practice in design of the vast majority of current clinical studies focusing on brain or SC neuroimaging. An alternative solution in future clinical study designs can be normalizing structural measurements for the body size or using body size as a confounding factor. In brain volume measurements, e.g., SIENAX^88^ or other kinds of normalization for the total intracranial volume may offer an effective normalization method that provides reproducible results independent of the body size. In the SC morphology, SIENAX^22^ or the dimension of pontomedullary junction^21^ have been implemented to normalize the CSA measurement. Yet, if possible, we conclude the body size matching provides a more optimal study design solution.

### Body size, neuroimaging and CNS (patho-)physiology

Body height had the highest impact on brain GM and SC WM morphology. Body height, higher cortical volume and improved cognitive ability appears to be phenotypically interlinked.^89^ The higher brain GM volumes in taller people may also explain their higher resistance to Alzheimer’s disease and other dementias.^90–92^ Gene expression could play a role here, as genetic variants that affect height also influence brain development and biological pathways that are involved in the development of Alzheimer’s disease.^90^

Although our data showed an insignificant interaction between body weight and CNS morphology after controlling for sex, body weight is known to influence CNS morphology and microstructure. Varying body weight showed WM and GM brain volume loss in patients with acute anorexia nervosa, and full WM volume and almost complete GM volume recoveries after the body weight had been regained.^93^ In the opposite body weight spectrum, obesity demonstrated lower intra-cortical myelination in regions involved in reward processing, attention, salience detection, and higher intra-cortical myelination in regions associated with somatosensory processing and inhibitory control.^94^ Increasing BMI changes cerebral WM microstructure assessed with DTI,^95^ but direction of DTI parameter trends in relation to body weight varies between studies.^96^ Although precise pathophysiological processes are not well known today, it is certain that obesity causes neuroinflammation, thus, alters brain microstructure and increases risks of neurodegenerative disorders such as Alzheimer’s disease and other types dementias.^97^ Our DTI and MTR data acquired in the current healthy population with low-to-moderate BMI may point to a borderline trend between homeostasis and mild microstructural changes related to the higher body weight. The negative correlation between body weight and MTR has also recently been reported in peripheral nerves and skeletal muscles.^35^ However, we cannot rule out the possibility of a transmit field (i.e., B1+) inhomogeneity-mediated bias in MTR. Although B1+ map was not measured for the cervical SC in our study, similar to what has been observed in the brain at 3T^98^, we expect both B1+ inhomogeneity and deviation to correlate with body weight positively, hence body transmit coil loading. Typically, an underflipping (i.e., reaching smaller than the desired flip angle) is more likely than an overflipping for small structures like the cervical SC in the body. MTR’s sensitivity to B1+ potentially exacerbates the effect of even a small degree of underflipping for the MT pulse at 3T.

Body height and spinal cord length are linearly dependent (r≈0.6).^99,100^ We show that even CSA-SC and CSA-WM are linearly dependent with body height. Thus, the magnitude of the correlation with body height would be even higher than observed for the CSA measurements if level-specific SC and SC WM volumes were analyzed. Although CSA values are level dependent,^24^ the impact of the C3-4 level selection on general study conclusions should remain minimal due to high intra-individual CSA correlation over segments.^101,102^ Different associations of CSA-GM and CSA-WM with other investigated variables, may affirm the necessity of further development of MRI protocols imaging SC GM in high contrast and detail.^103^

Recently, the correlation between CSA-SC at C2-3 level and body height, body weight, brain (WM/GM) volumes and thalamus volume were observed in 804 UK Biobank brain imaging database participants.^21^ Our current *spine-generic* database study complements the UK Biobank results and expands the knowledge that these observations are almost exclusively SC WM-related. Moreover, the current study identified more cerebral sub-regions involved than those investigated in the previous study. The lateral corticospinal tracts predominantly serving motor function are the major portion of the CSA-WM.^51^ Thus, its significant correlation with precentral gyrus thickness (primary motor cortex) seems logical from a neuroanatomical perspective. SC microstructure was also investigated and our exploratory approach via PCA clearly visualizes the body-SC-brain structural relationships.

The *spine-generic* database (*r20231212*) identifies 64 recruited subjects with the presence of degenerative cervical SC compression, with 2 of these even demonstrating radiological signs of myelopathy.^49^ These findings may represent a source of unexplained variance in our results, as compression and myelopathy are pathologies affecting CSA, DTI, and MTR measures.^64–66,104^ However, we showed in the **Supplementary Slides** that the impact of the compression on the correlation coefficient outcomes was minimal.

The observed negative correlation between age and cortical thickness and absence of correlation between body size and cortical thickness are in line with previous literature.^39,89,105–107^ The GM volume reduction in subcortical structures is less profound than in the cortical GM volume and thickness.^40,108,109^ Therefore, we may only detect low, insignificant trends in the age-related reductions of the subcortical structures due to an undersampled elderly population in our dataset. SC CSA-GM is also expected to decline with age,^37^ but we observed no such effects. Absent SC GM reduction might imply a false positive result due to the limited spatial resolution of the imaging methods, and the undersampled elderly population. It may also mean that the pathophysiological dynamics of SC GM reduction are slower than in the subcortical region. Yet, validating and concluding any of such statements require a rigorous re-test utilizing a dataset with a larger sample elderly population or longitudinal follow-up.

### Study limitations

Despite the relatively large sample size, there are still several limitations. First, we recruited healthy, predominantly young adults with average weight and low-to-moderate BMI. Therefore, the negative link between age and SC morphology, as observable in cohorts with greater age variability,^21,37,110^ was absent in our study. We found that body size impacts structural measurements more profoundly than age. However, this finding warrants further investigation, as the moderate age effects may be explained by the relatively narrow age range and younger cohort.^41^ However, concurrent study of 40-69-year-old adults also showed significant impact of body size on brain neuroimaging data.^46,87^ Head size was identified as an effective confounding factor minimizing the body size effects in brain structural measurements.^46,87^ The head size is not possible to measure precisely from the *spine-generic* database, because the images covering the brain were defaced. Thus, a significant portion of the image capturing head is missing in every scan. CSA-GM, SC DTI and SC MTR measurements demonstrated scanner-related variability, which needs to be addressed in multi-center data acquisition and analysis. Data of subjects with very low and high BMI may help to investigate the dependence of MTR and DTI measures on body weight. RF inhomogeneities need to be better mapped in future studies to avoid risks of biases in MTR outcomes. Comparison between SC and cerebral microstructure is impossible with the *spine-generic* database because the database does not contain images of brain microstructure. The current *spine-generic* database version does not allow assessing the impact of socioeconomic and race/ethnicity status on obtained MRI metrics.^111^ Relationships between spinal canal area,^99^ cervical cerebrospinal fluid area,^99^ and body size have not been investigated. Axial diffusivity (AD, i.e., another DTI metric) was not investigated. We expected that AD would provide similar results as observed for MD and RD due to expected high FA-MD-RD-AD intra-correlation levels. Therefore, we decided to shrink the variable space. Although the BMI correlated with several investigated neuroimaging variables,^23,35^ we reduced the variables to body height and weight only as BMI combines the two, and we expected similar findings. The cross-sectional study design limits testing of body size changes on the CNS over time.

## Conclusion

**(i)** We confirmed that “Future clinical research studies and trials utilizing neuroimaging should include body size as a potential confounding biological factor to avoid bias in clinical outcomes”. **(ii)** We hypothesized that “CSA of cervical SC WM and GM interacts with body size and morphology of distinct brain structures”, but after analysis we refine this to “CSA of cervical SC WM interacts with body height and morphology of distinct brain structures with a descending gradient from subcortical structures to cortical gray matter”. **(iii)** We hypothesized that “SC microstructure, as measured using MTR and DTI, interacts with body size”, but after analysis we refine this to “SC WM microstructure, as measured as MD and MTR, interacts with body weight, and more profoundly in dorsal columns than in lateral corticospinal tracts”. **(iv)** We confirmed our hypothesis that “Cerebral morphology interacts with body height more profoundly than with body weight and age”.

## Methods

### Structural MRI data

Signed informed consent was obtained from all participants under the compliance of the corresponding local ethics committee (more info in the *Scientific Data* paper^25^). The *spine-generic* protocol 3T MRI data were acquired once for each participant. Siemens scanners were used in 180 (67.41%) acquisitions, Philips scanners in 50 (18.72%) acquisitions and GE scanners in 37 (13.87%) acquisitions. 3D T1w scans were utilized to estimate cerebral volumes and cortical thicknesses. 3D T2w scans were utilized to assess the cross-sectional area (CSA) of the cervical spinal cord (SC). Axial T2*w scans were utilized to estimate the CSA of white (WM) and gray (GM) matter of the cervical SC. Diffusion weighted imaging was utilized to estimate diffusion tensor imaging (DTI) and the corresponding microstructural maps for the cervical SC. GRE-T1w, GRE-MT1, and GRE-MT0 scans were used to derive the magnetization transfer ratio (MTR) maps in the cervical SC. More detailed information about protocol settings and scanner subtypes can be found in the *spine-generic* protocol original papers.^24,25^

### Image analysis

The same image processing pipeline was employed here, utilizing the Spinal Cord Toolbox (SCT) version 6.1,^112^ as developed originally for the *spine-generic* protocol.^24,25^ CSA of the whole SC (CSA-SC) was computed and averaged from cervical C3-4 vertebral levels of the 3D T2w scan. CSA of WM and GM structures (CSA-WM, CSA-GM) were computed and averaged from cervical C3-4 levels of the axial T2*w scan. C3-4 levels were selected for CSA measurements since the T2*w imaging protocol had set the center of the field of view at the C3/4 disc and because C3-4 levels still contain the most sensory and motor fiber bundles. C3-4 average represents a robust representative morphological measurement as the CSA demonstrates high intra-individual correlation over segments,^101,102^ although the absolute CSA values inter-individually vary.^25^ All CSA measurements were measured in mm^2^ units. FA, MD, RD, and MTR were estimated from cervical C2-5 vertebral levels for WM, bilateral lateral corticospinal tracts, and bilateral dorsal columns utilizing the PAM50 atlas co-registration and weighted average techniques.^113^ The C2-5 segment range was selected for DTI and MTR averaging to guarantee the robustness of the tract-specific measurements with minimal partial volume effects.^113^

Brain volume was segmented and parceled at partial sub-structures from 3D T1w scans with FreeSurfer ver. 7.2.^114^ Volumes of brain (BrainVol), brain GM (BrainGMVol), cortical GM (CorticalGMVol), cortical WM (CorticalWMVol), subcortical GM (SubCortGMVol, including amygdala, caudate, hippocampus, nucleus accumbens, pallidum, putamen, thalamus, ventral diencephalon, and substancia nigra), thalamus (ThalamusVol), cerebellum (CerebellumVol), brainstem (BrainStemVol), precentral cortex GM (PrecentralGMVol) and postcentral cortex GM (PostcentralGMVol) were measured from the FreeSurfer segmentations in mm^3^ units. Cortical thickness (Cortical thickness), thickness of the precentral (PrecentralG Thickness), and postcentral gyrus (PostcentralG Thickness) were averaged across the left and right hemispheres as derived from the surface-based cortical parcellation. Precentral and postcentral cortices, motor and somatosensory cerebral centers, were investigated because the majority of the cervical spinal cord WM cross-section are the motor and somatosensory pathways.

### Exclusion of spinal cord and brain structural measurements

Spinal cord images were analyzed for all 267 participants. Cross-sectional area of SC (CSA-SC) was not estimated for 4 participants (listed in the category “csa_t2” in exclude.yml file, which contains the excluded subject ID and the verbal explanation of the exclusion; 1.50% of the dataset), CSA of WM and GM (CSA-WM and CSA-GM respectively) were not estimated for 4 different participants (category “csa_gm” in the exclude.yml file; 1.50%), DTI measurements were not estimated for 4 participants (categories “dti_fa”, “dti_md” and “dti_rd” in the exclude.yml file; 1.50%), and MTR measurements were not estimated for 5 participants (category “mtr” in the exclude.yml file; 1.87%). The exclude.yml file is available at: https://github.com/spine-generic/data-multi-subject/blob/r20231212/exclude.yml. The most common reasons for SC measurement exclusions were: (*i*) motion artifacts; (*ii*) subject repositioning during data acquisition; (*iii*) poor data quality; (*iv*) wrong field of view placement; or (*v*) not following required imaging parameters. The analysis excluded all CSA, DTI and MTR SC measurements for 1 additional subject (*sub-mniS05*; 0.37% of the dataset) due to severe degenerative cervical SC compression (maximal compression at C3/C4 level).

We analyzed brain images from 239 participants (89.51% of the dataset). We excluded T1w scans of 28 participants (10.49%) from the analysis because the images demonstrated field of view cut-offs (18 scans; 6.74%), defacing errors (5 scans; 1.87%), poor image contrast in superior cerebral regions (4 scans; 1.50%), and severe motion artifacts (1 scan; 0.37%). Excluded brain scans are listed in the exclude.yml file as the category “brain_t1”.

### Statistical analysis

Statistical analysis and figure visualization were implemented in the programming environment MATLAB R2021b (*Natick, USA*). Each variable or log(variable) was normalized into the space of the normal distribution and the Kruskall-Wallis test tested whether investigated variables meet conditions for Gaussian or log-Gaussian distribution (p<0.05). Between-group differences were tested with two-sample or paired t-tests.

Correlation analysis utilized Pearson (*r*) and Spearman (⍴) correlation coefficients, considering correlation to be significant if p_FWE_<0.05 (FWE - family wise error correction) after the Bonferroni multiple-comparison correction. Correlation coefficients were estimated for raw and normalized dataset values, where the manufacturer-specific average was subtracted from all SC quantitative measurements to minimize the effects of the previously reported inter-manufacturer variability in the spine-generic dataset^25^. For SC DTI and MTR correlation analysis, GE scanner raw values were excluded (i.e., 13.87% of the dataset) due to strong offsets compared to Siemens and Philips scanner values. SC qMRI measurements, FreeSurfer-based brain measurements, age, body height, and body weight were cross-correlated, and significant correlations (after the Bonferroni correction) were identified. The dataset was split into males and females and the correlation analysis was post-hoc repeated to address sex effects in the data demonstrating significant correlations. Due to the reduced sample size at half, the uncorrected p<0.05 was considered significant in the post-hoc analysis investigating the sex effects. The correlation analysis was also post-hoc repeated for SC measurements while excluding all 64 subjects with degenerative cervical SC compression (as identified in the *spine-generic* database; r20231212) to test for the compression effects on the study outcomes. The critical p_FWE_<0.05 remained here, although the dataset was reduced to 76% of its original size.

Manufacturer-specific average was subtracted from all SC structural measurements. Then, all multivariate data were normalized to mean=0 and standard deviation STD=1 for each examined variable. Such normalized data formed an input matrix for exploratory principal component analysis (PCA) optimized via singular value decomposition. Variables were visualized in the space of orthogonal principal components via biplot projections, and between-variable relationships were quantified and interpreted in the rotated principal space explaining the majority of the data variance.

Several linear regression models (Eqs. 1-6) were estimated for the SC and brain structural measurements (***y***) demonstrating significant correlation with age, body height and/or body weight respectively. Models’ coefficients of determination (*R*^2^) objectively assessed which demographic variable or set of demographic variables explained most of the demography-related variance in the SC and brain structure. Model utilizing simultaneous regression of body height and body weight was not utilized as body height and weight are strongly linearly dependent variables. The variable ***y****_0_* represents the model’s constant member, the *β* parameters are model regression coefficients (Eqs. 1-6). Categorical variable ***Sex*** was modeled as a vector of values 0.5 at positions of males and of values -0.5 at positions of females. Manufacturer-specific average was subtracted from all SC structural measurements before the regression analysis.

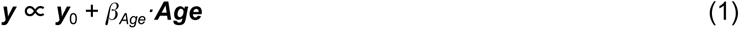

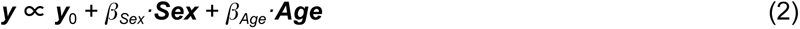

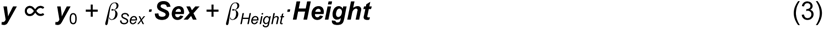

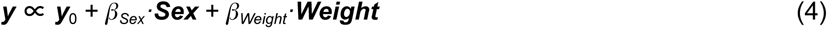

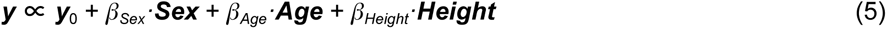

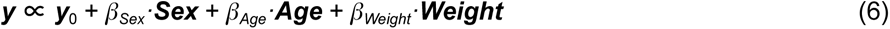

## Supporting information

Supplementary Slides

## Data availability

All raw data are publicly available at: https://github.com/spine-generic/data-multi-subject (utilized release ID: r20231212)

MRI imaging protocols for all optimized manufacturers and scanner types are publicly available at: https://github.com/spine-generic/protocols

Tables with SCT and Freesurfer measurements are available at: https://github.com/umn-milab/spine-generic-body-size-results (utilized release ID: r20240423)

## Computer code availability

Spinal Cord Toolbox is available at: https://github.com/spinalcordtoolbox/spinalcordtoolbox (utilized version: 6.1; git commit: git-master-c7a8072fd63a06a2775a74029c042833f0fce510)

FreeSurfer is available at: https://surfer.nmr.mgh.harvard.edu (utilized version: 7.2)

All computer code providing image and statistical analyses is available at: https://github.com/spine-generic/spine-generic (utilized release ID: height-weight-analysis)

## Acknowledgments

The Center for Magnetic Resonance Research (CMRR), Department of Radiology; the Center for Neurobehavioral Development (CNBD) at the Masonic Institute for the Developing Brain (MIDB), Department of Pediatrics; and the Minnesota Supercomputing Institute (MSI); all are part of the University of Minnesota and all provided lab space and computational support for this research project. The authors acknowledge the facilities, scientific and technical assistance of the National Imaging Facility, a National Collaborative Research Infrastructure Strategy (NCRIS) capability, at the Centre for Advanced Imaging, The University of Queensland. This work is based on experiments performed at the Swiss Center for Musculoskeletal Imaging, SCMI, Balgrist Campus AG, Zurich.

S Llufriu holds grants from the ISCIII [PI21/01189], AGAUR [021-SGR-01325] and research support from Bristol Myers Squibb. The research reported in this publication was also supported by the *National Natural Science Foundation of China* (82072010); the *Beijing Natural Science Foundation* [IS23108]; the *Ministry of Health, Czech Republic* - conceptual development of research organization [FNBr, 65269705]; the *Ministry of Education, Youth and Sports, Czech Republic* [LM2018129 Czech-BioImaging], part of the *Euro-BioImaging* (www.eurobioimaging.eu) *Advanced Light Microscopy and Medical Imaging Node* (Brno, Czech Republic); the *Max Planck Society* and the *European Research Council* [ERC StG 758974, European Union’s Horizon 2020 research and innovation programme 758974]; the *University of Pennsylvania* [MDBR-17-123-MPS - Million Dollar Bike Ride]; the *Instituto Salud Carlos III* - co-funded *European Union* [PI18/00823, PI22/01709]; the *Ministry of Health of the Czech Republi*c [NU22-04-00024]; the *European Union’s Horizon Europe* research and innovation programme under the Marie Skłodowska-Curie grant [101107932]; the *American Heart Association* [23CDA1054207]; the *Fondation Courtois*; the *Natural Sciences and Engineering Research Council of Canada* (NSERC), TransMedTech Institute, ICORD [RGPIN-2020–05242] and UBC; the *Craig H. Neilsen Foundation*; the *”la Caixa” Foundation* [ID 100010434; fellowship code LCF/BQ/PR22/11920010]; the *Wings For Life charity* [WFL-CH-19/20]; the *International Foundation for Research* [IRP-158]; the *Czech Health Research Council* [NV18-04-00159]; the *National Institute for Health Research Biomedical Research Centre at UCL and UCLH*; the *German Research Foundation* [TI 1110/1-1]; the *SpinalCure Australia*; the BRC [BRC1130/HEI/RS/11041]; the *European Union - NextGenerationEU* under the *Italian Ministry of University and Research (MUR) National Innovation Ecosystem* [ECS00000041 - VITALITY - CUP D73C22000840006]; the *Canada Research Chair in Quantitative Magnetic Resonance Imaging* [CRC-2020-00179]; the *Canadian Institute of Health Research* [PJT-190258]; the *Canada Foundation for Innovation* [32454, 34824]; the *Fonds de Recherche du Québec - Santé* [322736, 324636]; the *Natural Sciences and Engineering Research Council of Canada* [RGPIN-2019-07244]; the *Canada First Research Excellence Fund* (IVADO and TransMedTech), the *Courtois NeuroMod project*; the *Quebec BioImaging Network* [5886, 35450]; the *INSPIRED* (Spinal Research, UK; Wings for Life, Austria; Craig H. Neilsen Foundation, USA); the *Mila - Tech Transfer Funding Program*; and the *National Institutes of Health* (NIH) through the National Institute of Neurological Disorders and Stroke and the National Institute of Biomedical Imaging and Bioengineering [P41EB027061, P30NS076408, K23NS104211, L30NS108301, R01NS128478, R01NS133305, K01NS105160, K01EB030039, 5R01NS109114, K24NS126781, R61NS118651, R00EB016689, R01EB027779, R21EB031211, and R01NS109450]. The content is solely the responsibility of the authors and does not necessarily represent the official views of the NIH.

## Author contributions

RL initiated post-collection of missing demographic data, performed image analysis of spinal cord images and statistical analysis, designed figures and tables, and drafted the manuscript. MTB and ALP performed image analysis of brain images. EAO, SB, AF, JV and JCA provided data management and curation, and computer code integration into the *spine-generic* project. RL, SB, MA, EAO, NTA, LRB, RLB, MarBar, MarBat, CB, MDB, VC, AC, BDL, MDe, PLS, MDo, JD, AVD, FE, KRE, KSE, PF, JF, AF, MF, IF, CAMGW-K, GCG, GG, FeGi, FrGr, AH, P-GH, TH, MH, JMJ, KK, HK, MK, AK, J-WK, NK, HK, SK, YK, PeKu, PaKu, NDK, SK, MML, CoLa, CSWL, TL, YL, SL, SM, ARM, EM-H, LM, KPO’G, NP, DaPa, DePa, TBP, AP, FP, AR, MJR, RSS, GS, MS, ACS, AKS, SAS, ZAS, ES, YS, GT, AT, JV, DVDV, MCY, KAW II, NW, RGW, POW, JX, JCA, ChLe and IN collected data. JCA, ChLe and IN supervised RL, MTB and ALP in data analysis and substantially revised the first manuscript draft. *<u>All authors</u>* designed and conceptualized the study, read and revised the manuscript.

