## Supplementary Materials for "Body size interacts with the structure of the central nervous system: A multi-center in vivo neuroimaging study"

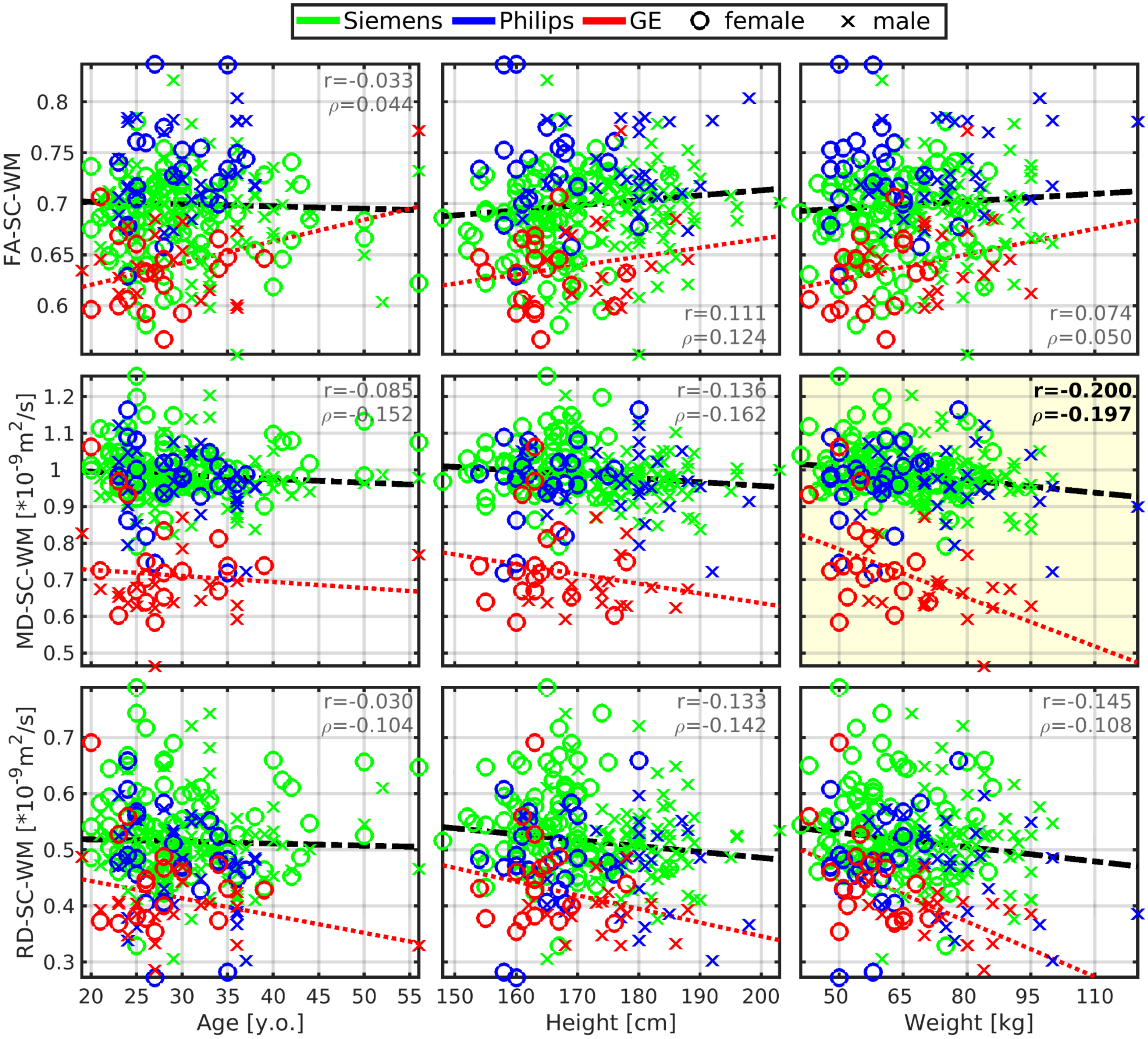

**Figure S1: Mean diffusivity in spinal cord (SC) white matter (WM) correlates to body weight.**
_Abbreviations:_ WM - white matter; SC - spinal cord; FA - fractional anisotropy; MD - mean diffusivity; RD - radial diffusivity; r - Pearson correlation coefficient; ρ - Spearman correlation coefficient. All spinal cord measurements were averaged from cervical C2-5 levels. Black dashed regression lines were estimated from the Siemens and Philips scanners' data points. Red dotted regression lines were estimated from the GE scanner's data points. Plots with statistically significant correlation ($p_{FWE}<0.05$) are highlighted with yellow background, and corresponding r and ρ values are highlighted with black bold font.

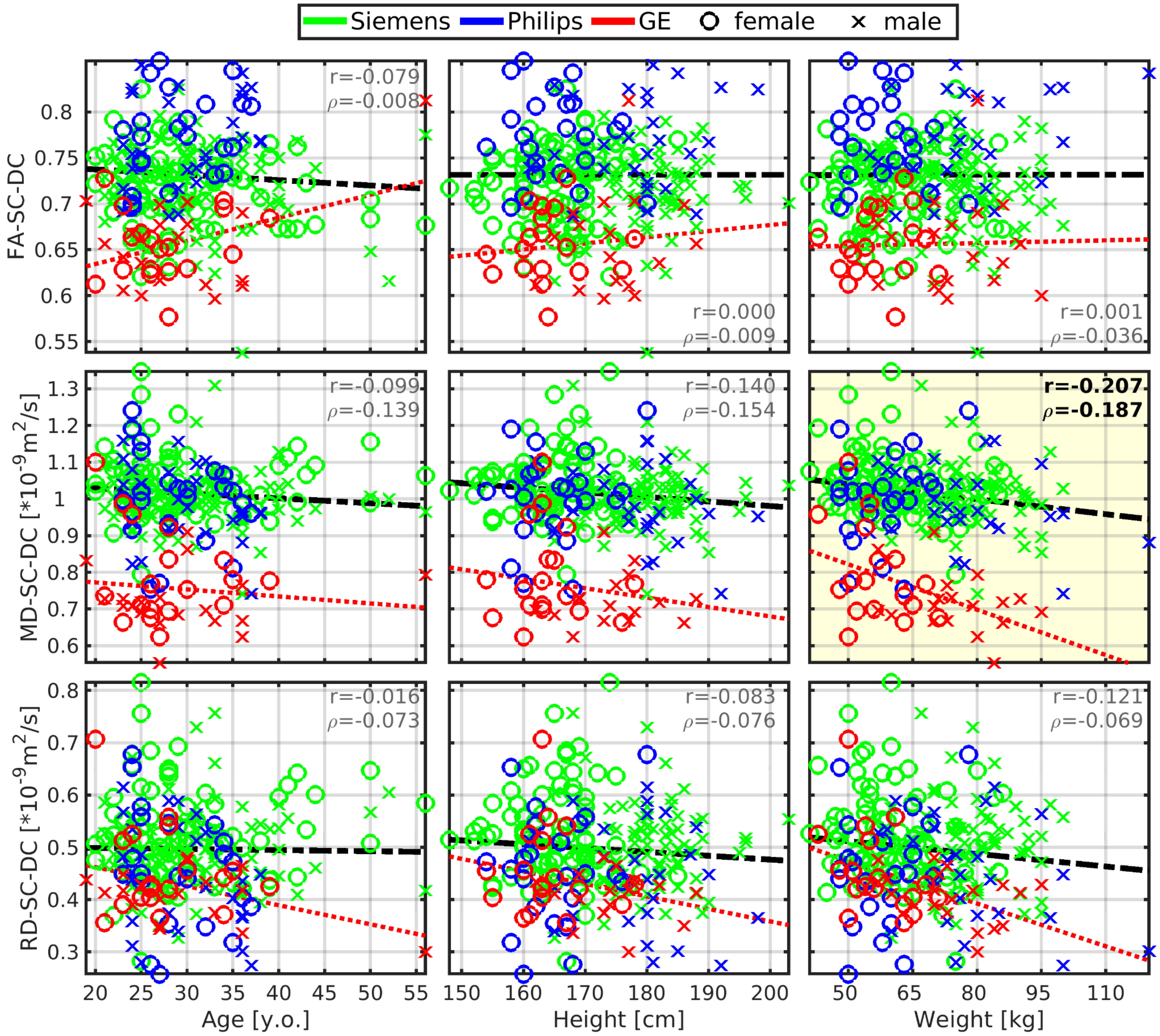

**Figure S2: Mean diffusivity in spinal cord (SC) dorsal columns (DC) is correlated to body weight.**

*Abbreviations:* DC - dorsal columns; SC - spinal cord; FA - fractional anisotropy; MD - mean diffusivity; RD - radial diffusivity; r - Pearson correlation coefficient; ρ - Spearman correlation coefficient. All spinal cord measurements were averaged from cervical C2-5 levels. Black dashed regression lines were estimated from the Siemens and Philips scanners' data points. Red dotted regression lines were estimated from the GE scanner's data points. Plots with statistically significant correlation ($p_{FWE}$<0.05) are highlighted with yellow background, and corresponding r and ρ values are highlighted with black bold font.

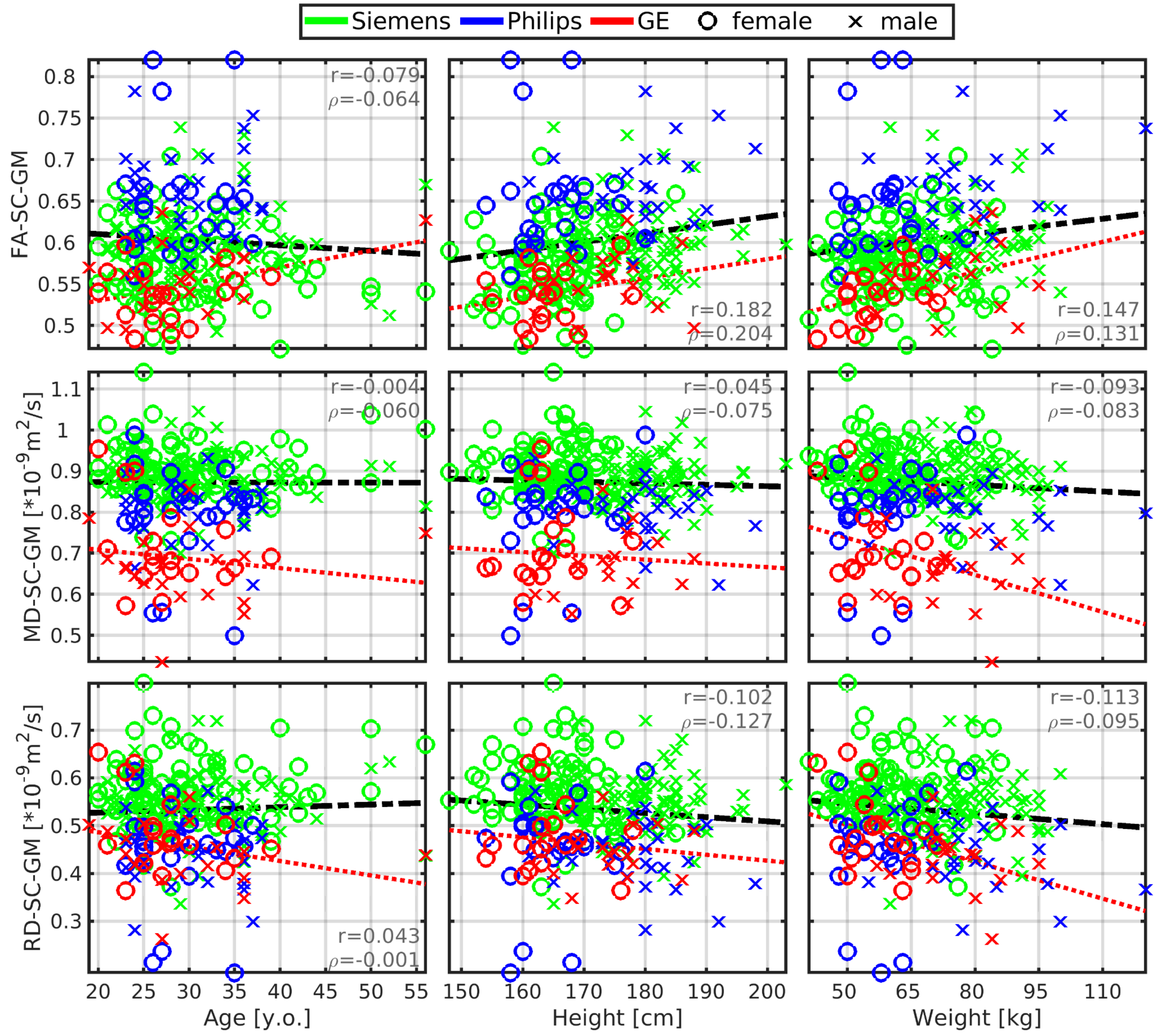

**Figure S3: No correlation between diffusion tensor imaging in spinal cord (SC) gray matter (GM) and body size or age respectively.**

*Abbreviations:* GM - gray matter; SC - spinal cord; FA - fractional anisotropy; MD - mean diffusivity; RD - radial diffusivity; r - Pearson correlation coefficient; ρ - Spearman correlation coefficient. All spinal cord measurements were averaged from cervical C2-5 levels. Black dashed regression lines were estimated from the Siemens and Philips scanners' data points. Red dotted regression lines were estimated from the GE scanner's data points. No plotted correlations are significant ($p_{FWE}$<0.05).

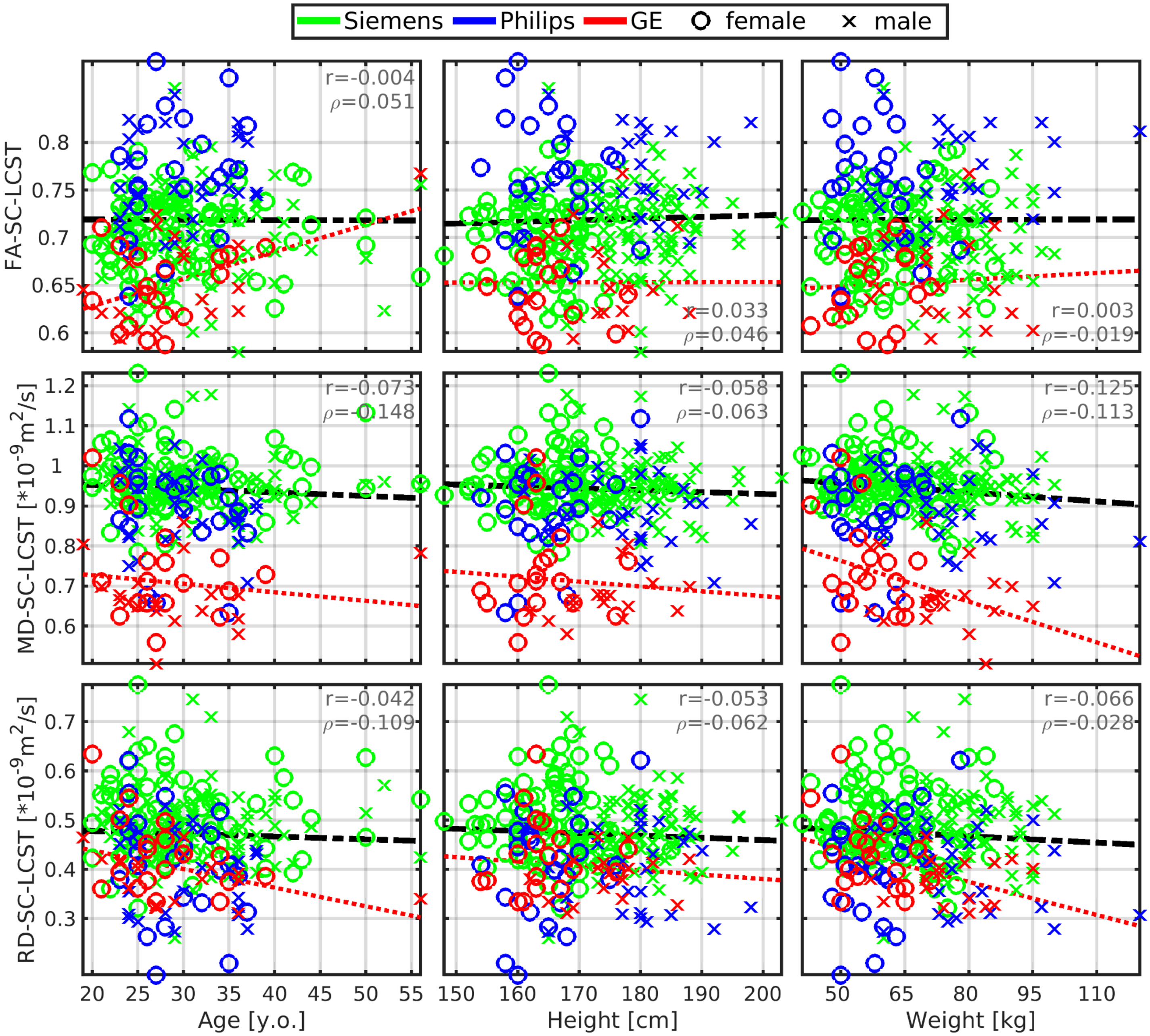

**Figure S4: No correlation between diffusion tensor imaging in spinal cord lateral corticospinal tracts (LCST) and body size or age respectively.**

_Abbreviations:_ LCST - lateral corticospinal tracts; SC - spinal cord; FA - fractional anisotropy; MD - mean diffusivity; RD - radial diffusivity; r - Pearson correlation coefficient; ρ - Spearman correlation coefficient. All spinal cord measurements were averaged from cervical C2-5 levels. Black dashed regression lines were estimated from the Siemens and Philips scanners' data points. Red dotted regression lines were estimated from the GE scanner's data points. No plotted correlations are significant ($p_{FWE}<0.05$).

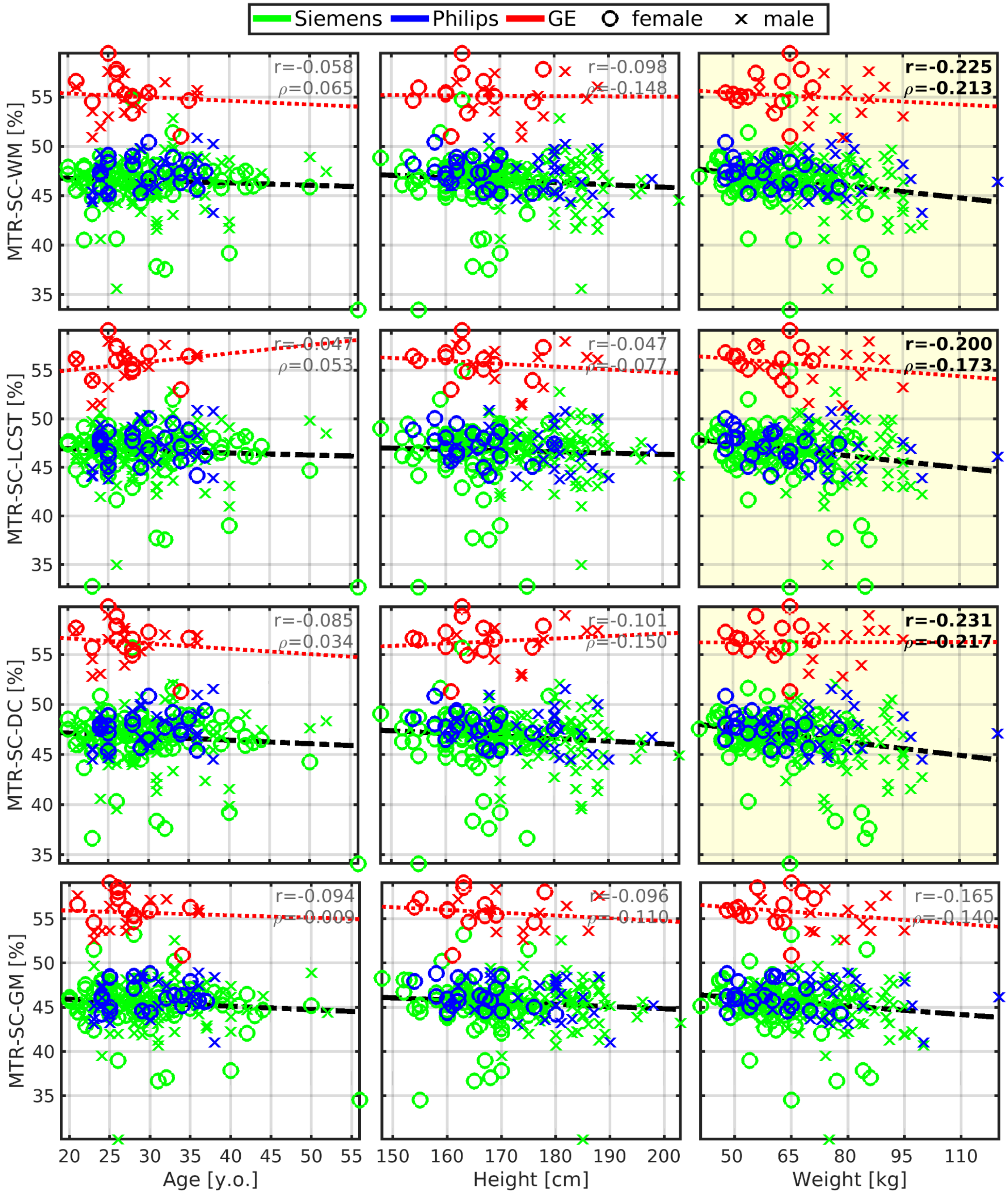

**Figure S5: Spinal cord magnetization transfer ratio imaging is correlated to body weight.**

*Abbreviations:* WM - white matter; SC - spinal cord; LCST - lateral corticospinal tracts; DC - dorsal columns; GM - gray matter; MTR - magnetization transfer ratio; r - Pearson correlation coefficient; ρ - Spearman correlation coefficient. All spinal cord measurements were averaged from cervical C2-5 levels. Black dashed regression lines were estimated from the Siemens and Philips scanners' data points. Red dotted regression lines were estimated from the GE scanner's data points. Plots with statistically significant correlation ($p_{FWE}$<0.05) are highlighted with yellow background, and corresponding r and ρ values are highlighted with black bold font.

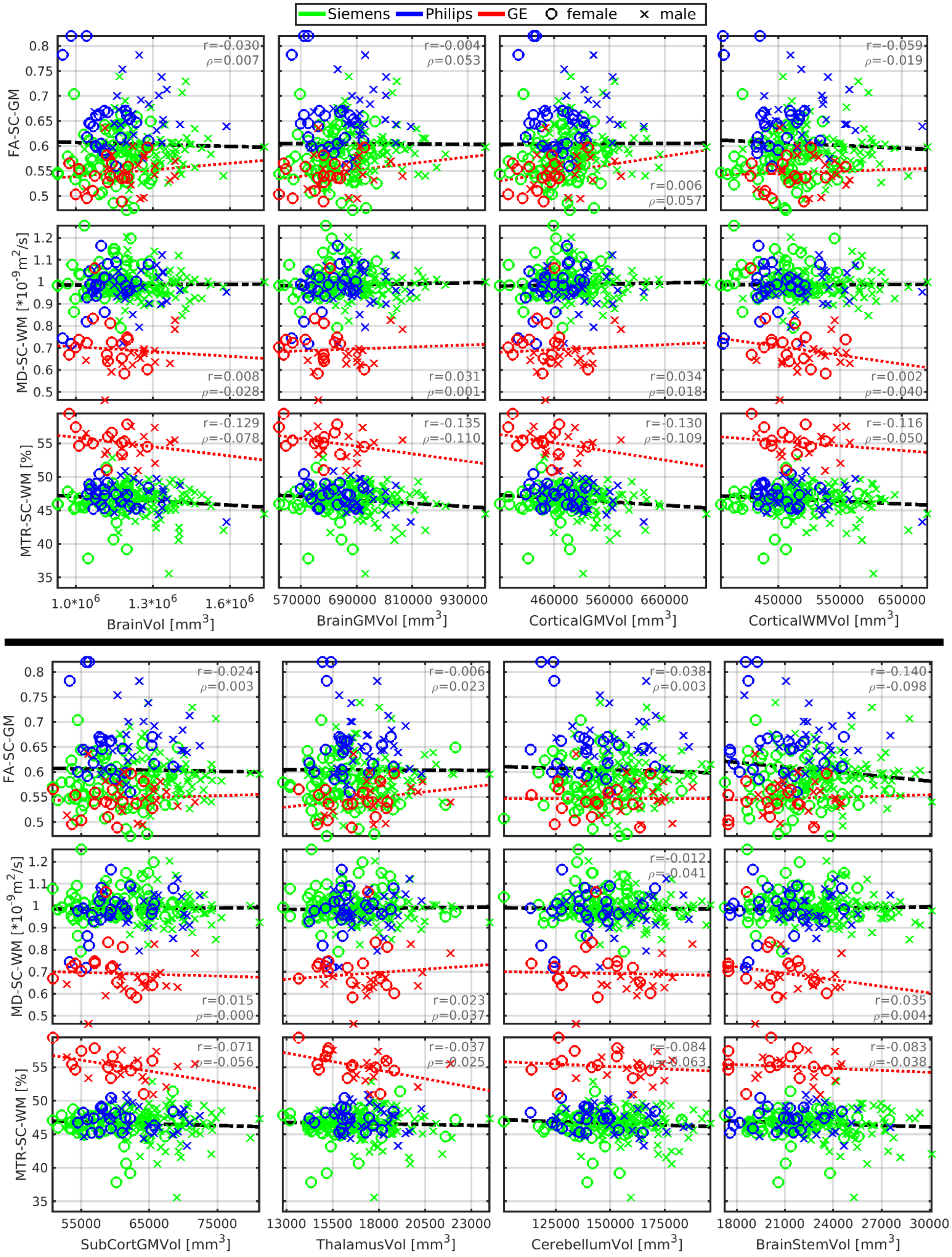

**Figure S6: No links between cerebral macrostructure (x-axis) and spinal cord microstructure (y-axis).** *Abbreviations:* WM - white matter; SC - spinal cord; FA - fractional anisotropy; MD - mean diffusivity; MTR - magnetization transfer ratio; GM - gray matter; Vol - volume; SubCort - subcortical; r - Pearson correlation coefficient; ρ - Spearman correlation coefficient. Regression lines (i.e., the dashed black lines) were estimated from all available data points. No plotted correlations are significant ($p_{FWE}$<0.05).

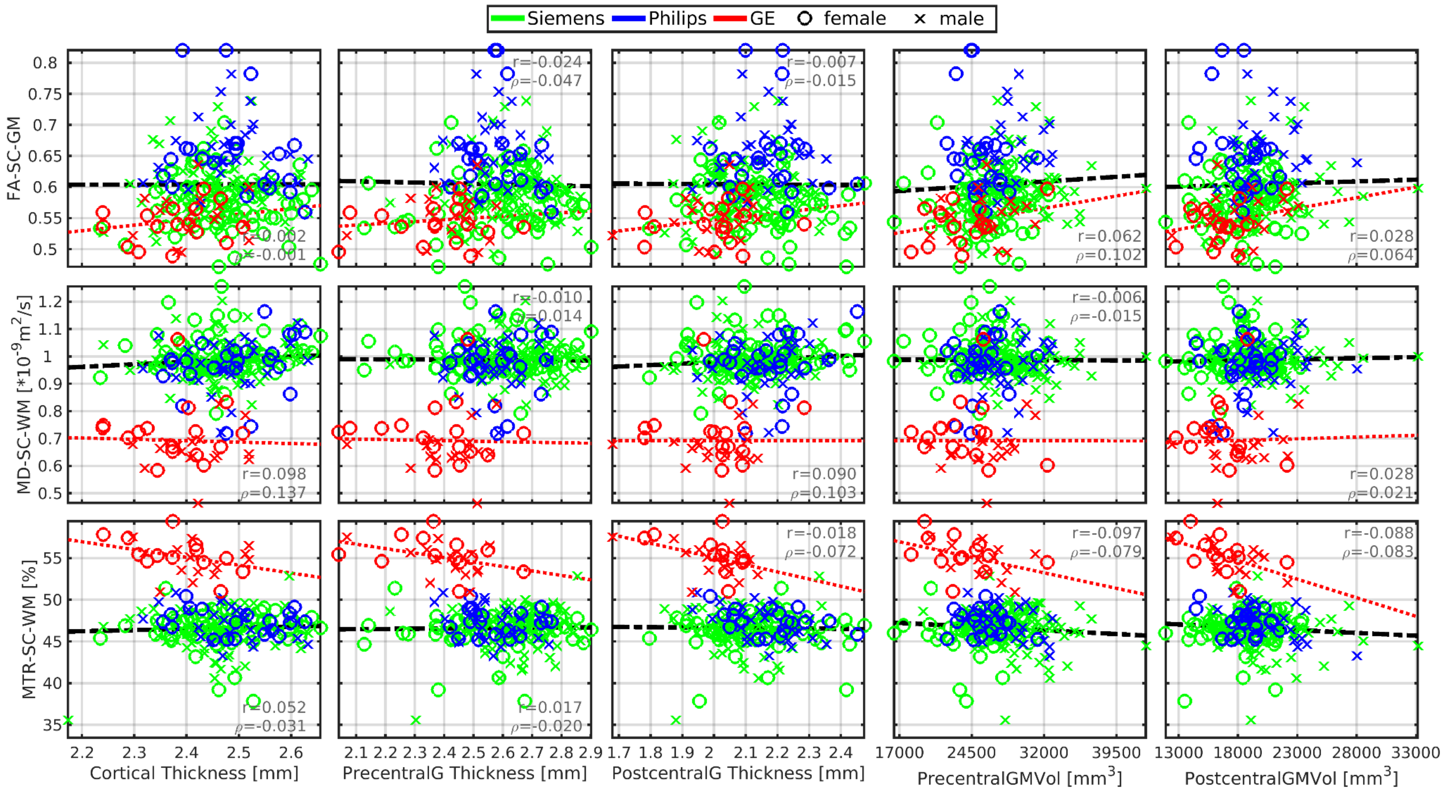

**Figure S7: No links between cortical macrostructure (x-axis) and spinal cord microstructure (y-axis).**
*Abbreviations:* WM - white matter; SC - spinal cord; FA - fractional anisotropy; MD - mean diffusivity; MTR - magnetization transfer ratio; GM - gray matter; Vol - volume; r - Pearson correlation coefficient; ρ - Spearman correlation coefficient. Regression lines (i.e., the dashed black lines) were estimated from all available data points. No plotted correlations are significant ($p_{FWE}$<0.05).

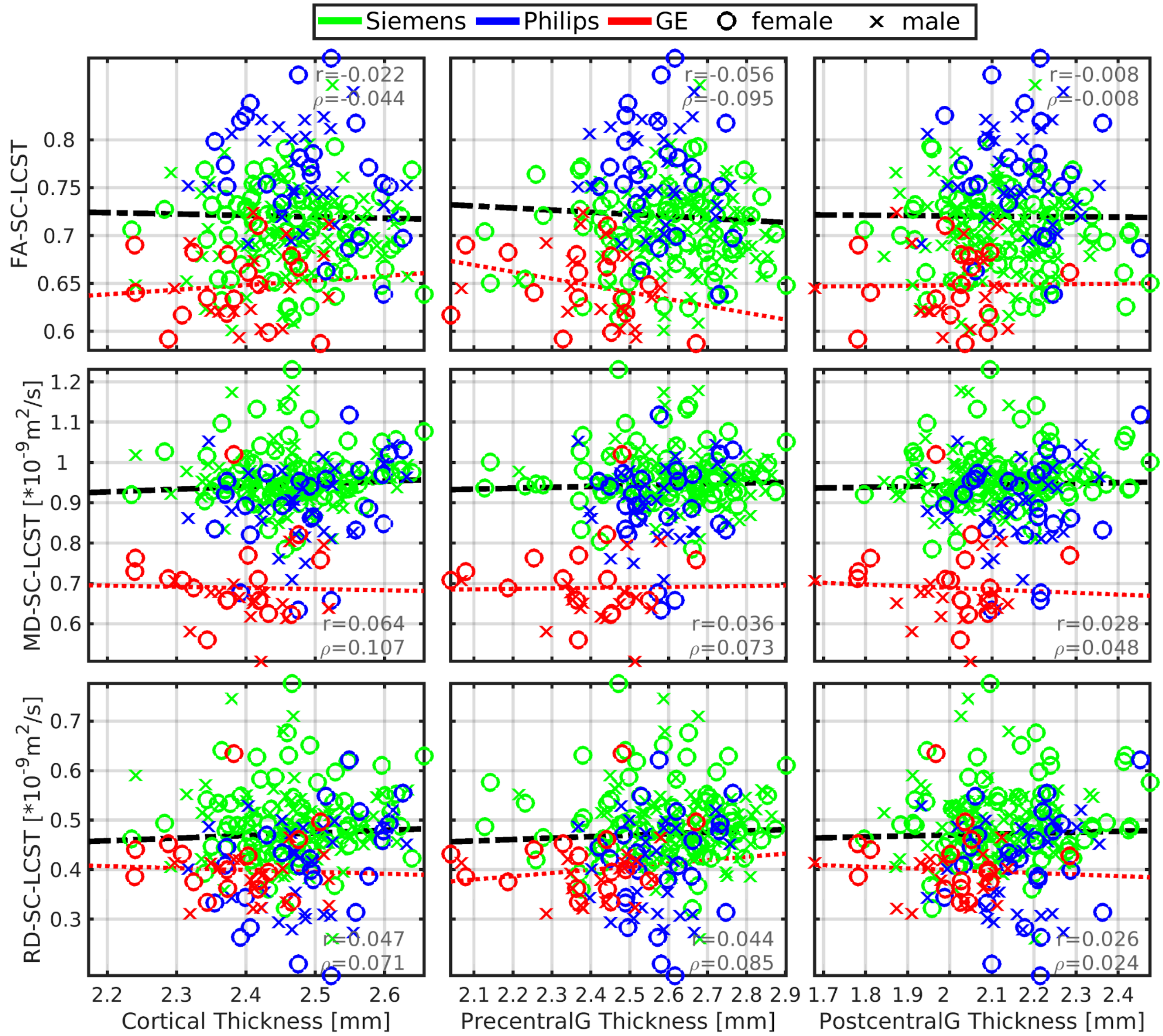

**Figure S8: No links between cortical thicknesses (x-axis) and spinal cord diffusion tensor imaging of bilateral lateral corticospinal tracts (y-axis).**

Abbreviations: LCST - lateral corticospinal tracts; SC - spinal cord; FA - fractional anisotropy; MD - mean diffusivity; RD - radial diffusivity; r - Pearson correlation coefficient; ρ - Spearman correlation coefficient. All spinal cord measurements were averaged from cervical C2-5 levels. Black dashed regression lines were estimated from the Siemens and Philips scanners' data points. Red dotted regression lines were estimated from the GE scanner's data points.No plotted correlations are significant ($p_{FWE}<0.05$).

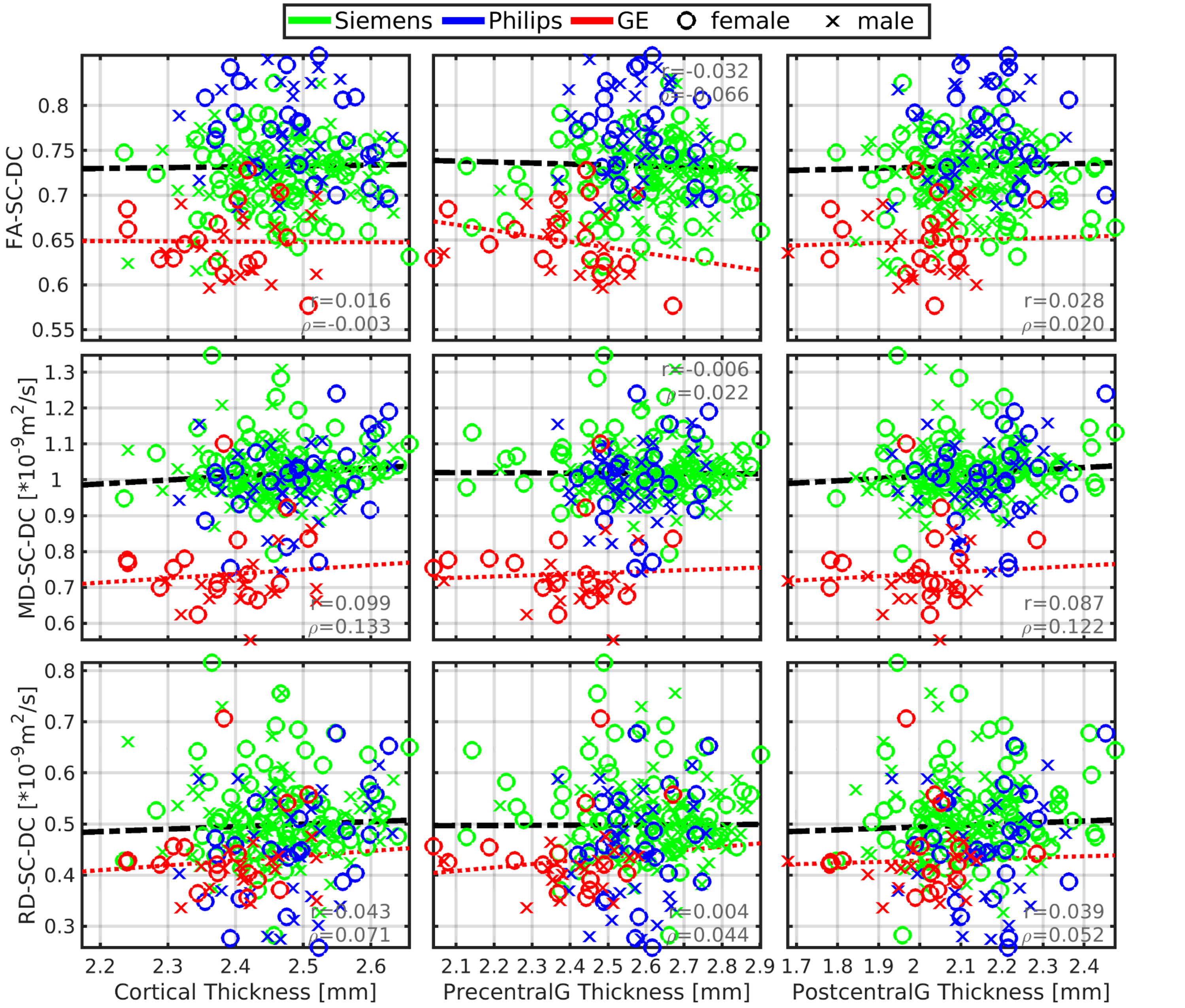

**Figure S9: No links between cortical thicknesses (x-axis) and spinal cord diffusion tensor imaging of bilateral dorsal columns (y-axis).**

_Abbreviations:_ DC - dorsal columns; SC - spinal cord; FA - fractional anisotropy; MD - mean diffusivity; RD - radial diffusivity; r - Pearson correlation coefficient; ρ - Spearman correlation coefficient. All spinal cord measurements were averaged from cervical C2-5 levels. Black dashed regression lines were estimated from the Siemens and Philips scanners' data points. Red dotted regression lines were estimated from the GE scanner's data points. No plotted correlations are significant ($p_{FWE}$<0.05).

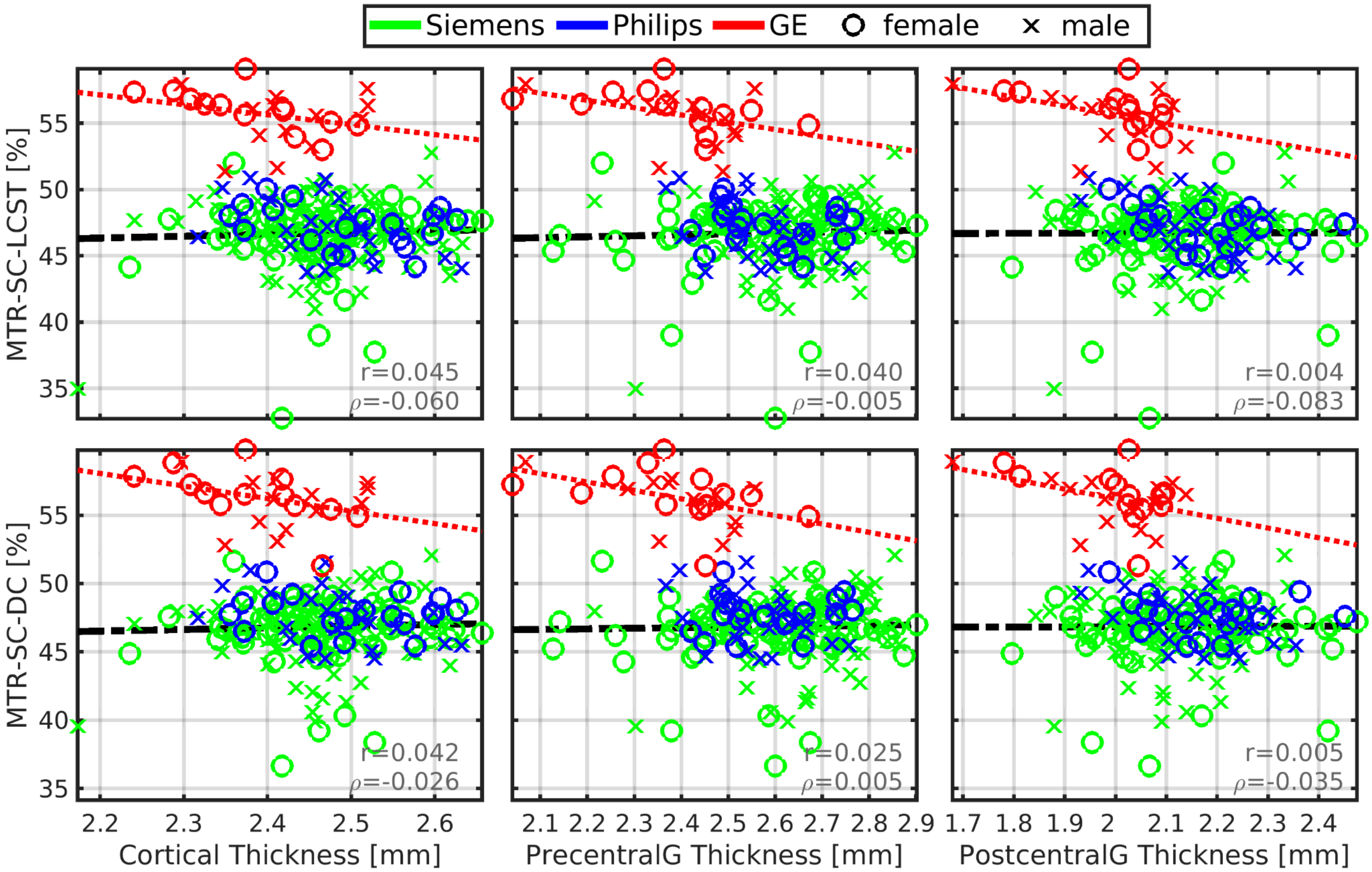

**Figure S10: No links between cortical thicknesses (x-axis) and magnetization transfer ratio of bilateral lateral corticospinal tracts and bilateral dorsal columns (y-axis).**

_Abbreviations:_ SC - spinal cord; LCST - lateral corticospinal tracts; DC - dorsal columns; MTR - magnetization transfer ratio; r - Pearson correlation coefficient; ρ - Spearman correlation coefficient. All spinal cord measurements were averaged from cervical C2-5 levels. Black dashed regression lines were estimated from the Siemens and Philips scanners' data points. Red dotted regression lines were estimated from the GE scanner's data points. No plotted correlations are significant ($p_{FWE}<0.05$).

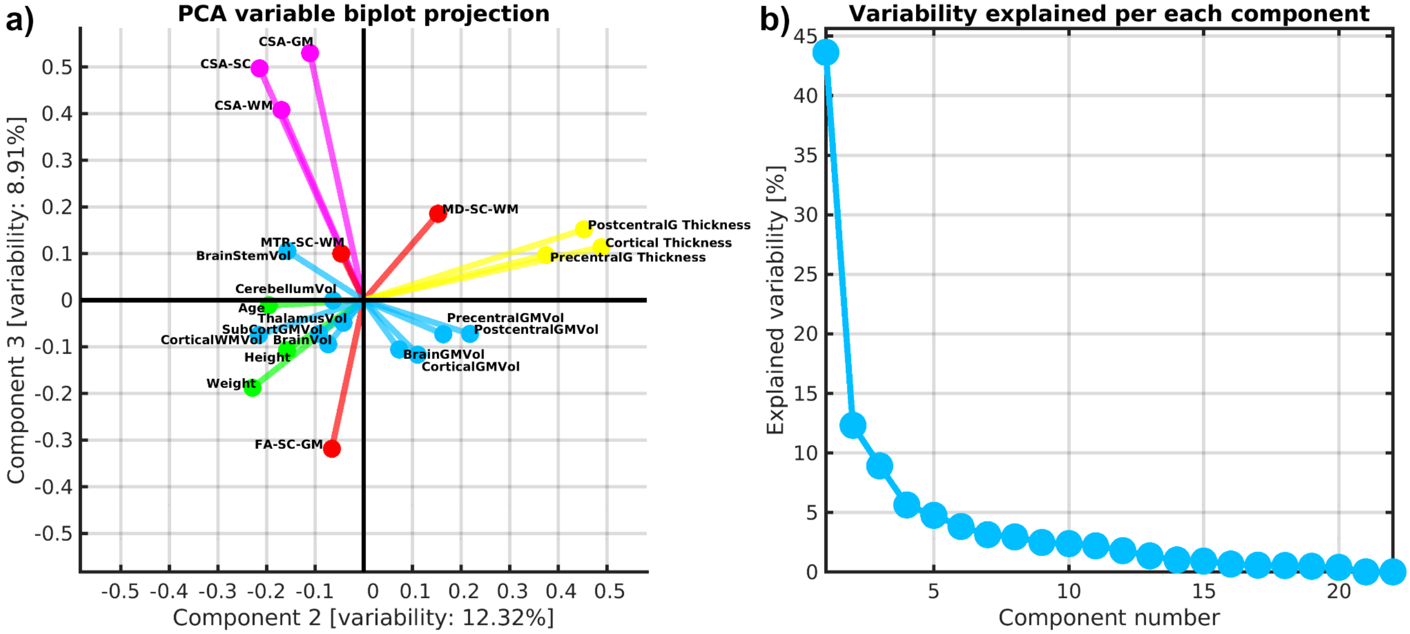

**Figure S11: Biplot projection of 2nd and 3rd principal components (a) and explained data variability per principal component (b).**

**Supplementary Table 1: Full table of Pearson correlation coefficients (r) between body size, age, spinal cord structure, and brain structure, and post-hoc sex-effects in the correlation analysis.**

*Abbreviations:* CSA - cross-sectional area; SC - spinal cord; WM - white matter; GM - gray matter; Vol - volume.

The correlation p-values (p) are uncorrected. Table 2 in the main paper is a subset of this supplementary table. CSA was measured as averages between C3-C4 segments. DTI and MTR were calculated as averages between C2-C5 segments. The column denoted "Original values" reports correlation coefficients for raw measurements with no normalization procedure prior to the correlation analysis. The column denoted "Manufacturer-specific normalized SC measures" reports correlation coefficients for SC structural measurements, which were normalized to zero mean for each scanner manufacturer before correlation analysis. Empty cells in the right half of the table represent combinations where no updated correlation coefficients were measured, because the utilized normalization of SC structural measurements had no effect on these correlation coefficients. Brain structural measurements were not considered necessary to normalize as we did not observe strong scanner-related effects in brain macrostructural measurements.

| | **Original values** | | | | | | **Manufacturer-specific normalized values** | | | | | |
|---|---|---|---|---|---|---|---|---|---|---|---|---|
| | **All** | | **Female** | | **Male** | | **All** | | **Female** | | **Male** | |
| **Correlation pair** | **r** | **p** | **r** | **p** | **r** | **p** | **r** | **p** | **r** | **p** | **r** | **p** |
| Age vs CSA-SC | 0.047 | 4.5E-1 | 0.031 | 7.3E-1 | 0.038 | 6.6E-1 | 0.029 | 6.5E-1 | 0.004 | 9.7E-1 | 0.024 | 7.8E-1 |
| Age vs CSA-WM | -0.001 | 9.8E-1 | -0.015 | 8.7E-1 | -0.043 | 6.2E-1 | -0.026 | 6.7E-1 | -0.049 | 5.9E-1 | -0.064 | 4.5E-1 |
| Age vs CSA-GM | 0.038 | 5.4E-1 | -0.003 | 9.7E-1 | 0.070 | 4.1E-1 | -0.001 | 9.9E-1 | -0.053 | 5.6E-1 | 0.039 | 6.5E-1 |
| Height vs CSA-SC | 0.355 | **5.0E-9** | 0.323 | **2.9E-4** | 0.230 | **7.3E-3** | 0.344 | **1.5E-8** | 0.319 | **3.5E-4** | 0.205 | **1.7E-2** |
| Height vs CSA-WM | 0.437 | **1.7E-13** | 0.295 | **9.9E-4** | 0.303 | **3.2E-4** | 0.422 | **1.4E-12** | 0.285 | **1.4E-3** | 0.268 | **1.5E-3** |
| Height vs CSA-GM | 0.070 | 2.6E-1 | 0.148 | 1.0E-1 | 0.052 | 5.4E-1 | 0.092 | 1.4E-1 | 0.213 | **1.9E-2** | 0.026 | 7.7E-1 |
| Weight vs CSA-SC | 0.261 | **2.2E-5** | 0.140 | 1.2E-1 | 0.154 | 7.3E-2 | 0.256 | **3.1E-5** | 0.108 | 2.4E-1 | 0.159 | 6.3E-2 |
| Weight vs CSA-WM | 0.274 | **7.3E-6** | 0.100 | 2.7E-1 | 0.084 | 3.2E-1 | 0.266 | **1.4E-5** | 0.071 | 4.3E-1 | 0.077 | 3.7E-1 |
| Weight vs CSA-GM | -0.021 | 7.3E-1 | 0.056 | 5.4E-1 | -0.089 | 3.0E-1 | -0.020 | 7.5E-1 | 0.002 | 9.8E-1 | -0.075 | 3.8E-1 |
| Age vs FA-WM | -0.033 | 6.2E-1 | 0.027 | 7.8E-1 | -0.108 | 2.4E-1 | 0.043 | 4.9E-1 | 0.057 | 5.3E-1 | 0.011 | 9.0E-1 |
| Age vs MD-WM | -0.085 | 2.0E-1 | -0.024 | 8.1E-1 | -0.124 | 1.8E-1 | -0.093 | 1.3E-1 | -0.079 | 3.8E-1 | -0.077 | 3.7E-1 |
| Age vs RD-WM | -0.030 | 6.5E-1 | -0.022 | 8.2E-1 | -0.014 | 8.8E-1 | -0.080 | 2.0E-1 | -0.070 | 4.4E-1 | -0.060 | 4.8E-1 |
| Height vs FA-WM | 0.111 | 9.8E-2 | 0.064 | 5.1E-1 | 0.040 | 6.6E-1 | 0.132 | **3.4E-2** | 0.060 | 5.1E-1 | 0.068 | 4.3E-1 |
| Height vs MD-WM | -0.136 | **4.1E-2** | 0.126 | 2.0E-1 | -0.125 | 1.8E-1 | -0.147 | **1.8E-2** | 0.082 | 3.7E-1 | -0.088 | 3.1E-1 |
| Height vs RD-WM | -0.133 | **4.7E-2** | 0.041 | 6.8E-1 | -0.074 | 4.3E-1 | -0.156 | **1.2E-2** | 0.022 | 8.1E-1 | -0.073 | 4.0E-1 |
| Weight vs FA-WM | 0.074 | 2.7E-1 | -0.013 | 9.0E-1 | 0.031 | 7.4E-1 | 0.103 | 9.8E-2 | 0.054 | 5.5E-1 | 0.018 | 8.3E-1 |
| Weight vs MD-WM | -0.200 | **2.6E-3** | -0.022 | 8.2E-1 | -0.191 | **3.8E-2** | -0.252 | **3.8E-5** | -0.108 | 2.3E-1 | -0.206 | **1.6E-2** |
| Weight vs RD-WM | -0.145 | **3.0E-2** | 0.007 | 9.4E-1 | -0.108 | 2.4E-1 | -0.203 | **9.6E-4** | -0.080 | 3.8E-1 | -0.124 | 1.5E-1 |
| Age vs MTR-WM | -0.058 | 4.0E-1 | -0.233 | **2.0E-2** | 0.125 | 1.9E-1 | -0.053 | 4.2E-1 | -0.255 | **6.6E-3** | 0.157 | 8.0E-2 |
| Height vs MTR-WM | -0.098 | 1.6E-1 | -0.090 | 3.8E-1 | -0.189 | **4.6E-2** | -0.091 | 1.6E-1 | -0.065 | 5.0E-1 | -0.151 | 9.4E-2 |
| Weight vs MTR-WM | -0.225 | **1.0E-3** | -0.374 | **1.4E-4** | -0.221 | **1.9E-2** | -0.221 | **6.1E-4** | -0.331 | **3.6E-4** | -0.217 | **1.5E-2** |
| BrainVol vs CSA-SC | 0.419 | **3.2E-11** | 0.337 | **3.7E-4** | 0.368 | **2.8E-5** | 0.393 | **6.1E-10** | 0.311 | **1.0E-3** | 0.327 | **2.2E-4** |
| BrainVol vs CSA-WM | 0.524 | **9.7E-18** | 0.392 | **2.9E-5** | 0.459 | **7.5E-8** | 0.507 | **1.4E-16** | 0.379 | **5.7E-5** | 0.426 | **7.3E-7** |
| BrainVol vs CSA-GM | 0.221 | **7.0E-4** | 0.250 | **9.4E-3** | 0.298 | **7.3E-4** | 0.186 | **4.4E-3** | 0.219 | **2.3E-2** | 0.254 | **4.3E-3** |
| BrainGMVol vs CSA-SC | 0.357 | **2.4E-8** | 0.265 | **5.6E-3** | 0.287 | **1.3E-3** | 0.324 | **4.9E-7** | 0.225 | **1.9E-2** | 0.242 | **7.0E-3** |
| BrainGMVol vs CSA-WM | 0.481 | **7.9E-15** | 0.341 | **3.3E-4** | 0.406 | **2.7E-6** | 0.445 | **1.2E-12** | 0.286 | **2.8E-3** | 0.359 | **3.9E-5** |
| BrainGMVol vs CSA-GM | 0.159 | **1.5E-2** | 0.147 | 1.3E-1 | 0.228 | **1.1E-2** | 0.128 | 5.1E-2 | 0.133 | 1.7E-1 | 0.182 | **4.3E-2** |
| CorticalGMVol vs CSA-SC | 0.320 | **7.0E-7** | 0.195 | **4.3E-2** | 0.262 | **3.4E-3** | 0.290 | **7.2E-6** | 0.159 | 1.0E-1 | 0.225 | **1.2E-2** |

| | | | | | | | | | | | | |
|---|---|---|---|---|---|---|---|---|---|---|---|---|
| CorticalGMVol vs CSA-WM | 0.449 | **6.7E-13** | 0.283 | **3.2E-3** | 0.390 | **7.0E-6** | 0.412 | **6.4E-11** | 0.227 | **1.9E-2** | 0.346 | **7.6E-5** |
| CorticalGMVol vs CSA-GM | 0.124 | 5.9E-2 | 0.102 | 3.0E-1 | 0.177 | **4.8E-2** | 0.106 | 1.1E-1 | 0.093 | 3.4E-1 | 0.154 | 8.7E-2 |
| CorticalWMVol vs CSA-SC | 0.432 | **6.2E-12** | 0.291 | **2.3E-3** | 0.422 | **1.1E-6** | 0.411 | **8.0E-11** | 0.280 | **3.3E-3** | 0.384 | **1.2E-5** |
| CorticalWMVol vs CSA-WM | 0.503 | **2.7E-16** | 0.315 | **9.5E-4** | 0.471 | **2.9E-8** | 0.501 | **3.6E-16** | 0.339 | **3.6E-4** | 0.447 | **1.8E-7** |
| CorticalWMVol vs CSA-GM | 0.268 | **3.6E-5** | 0.275 | **4.1E-3** | 0.355 | **4.9E-5** | 0.217 | **8.6E-4** | 0.215 | **2.6E-2** | 0.301 | **6.6E-4** |
| SubCortGMVol vs CSA-SC | 0.417 | **3.8E-11** | 0.207 | **3.2E-2** | 0.449 | **1.8E-7** | 0.388 | **9.7E-10** | 0.198 | **4.0E-2** | 0.392 | **7.4E-6** |
| SubCortGMVol vs CSA-WM | 0.511 | **7.8E-17** | 0.257 | **7.5E-3** | 0.524 | **3.4E-10** | 0.488 | **2.9E-15** | 0.248 | **1.0E-2** | 0.474 | **2.4E-8** |
| SubCortGMVol vs CSA-GM | 0.217 | **9.0E-4** | 0.093 | 3.4E-1 | 0.392 | **6.1E-6** | 0.197 | **2.6E-3** | 0.119 | 2.2E-1 | 0.343 | **8.9E-5** |
| ThalamusVol vs CSA-SC | 0.335 | **1.8E-7** | 0.176 | 6.8E-2 | 0.322 | **2.8E-4** | 0.338 | **1.4E-7** | 0.201 | **3.7E-2** | 0.308 | **5.3E-4** |
| ThalamusVol vs CSA-WM | 0.431 | **6.3E-12** | 0.244 | **1.1E-2** | 0.398 | **4.4E-6** | 0.451 | **5.0E-13** | 0.279 | **3.6E-3** | 0.408 | **2.4E-6** |
| ThalamusVol vs CSA-GM | 0.166 | **1.1E-2** | 0.058 | 5.5E-1 | 0.297 | **7.7E-4** | 0.211 | **1.2E-3** | 0.159 | 1.0E-1 | 0.316 | **3.3E-4** |
| CerebellumVol vs CSA-SC | 0.356 | **2.5E-8** | 0.508 | **2.0E-8** | 0.109 | 2.3E-1 | 0.342 | **9.9E-8** | 0.493 | **6.0E-8** | 0.086 | 3.4E-1 |
| CerebellumVol vs CSA-WM | 0.433 | **4.9E-12** | 0.501 | **3.8E-8** | 0.179 | **4.6E-2** | 0.441 | **1.8E-12** | 0.508 | **2.4E-8** | 0.186 | **3.8E-2** |
| CerebellumVol vs CSA-GM | 0.200 | **2.3E-3** | 0.325 | **6.4E-4** | 0.183 | **4.1E-2** | 0.187 | **4.3E-3** | 0.374 | **7.3E-5** | 0.121 | 1.8E-1 |
| BrainStemVol vs CSA-SC | 0.575 | **1.1E-21** | 0.622 | **6.5E-13** | 0.492 | **7.4E-9** | 0.519 | **2.5E-17** | 0.573 | **9.1E-11** | 0.409 | **2.6E-6** |
| BrainStemVol vs CSA-WM | 0.640 | **4.0E-28** | 0.624 | **6.8E-13** | 0.547 | **4.0E-11** | 0.590 | **3.8E-23** | 0.580 | **6.0E-11** | 0.467 | **4.1E-8** |
| BrainStemVol vs CSA-GM | 0.390 | **7.2E-10** | 0.444 | **1.6E-6** | 0.480 | **1.5E-8** | 0.335 | **1.8E-7** | 0.440 | **2.1E-6** | 0.375 | **1.6E-5** |
| PrecentralGMVol vs CSA-SC | 0.307 | **1.9E-6** | 0.159 | 1.0E-1 | 0.273 | **2.1E-3** | 0.275 | **2.2E-5** | 0.132 | 1.7E-1 | 0.228 | **1.1E-2** |
| PrecentralGMVol vs CSA-WM | 0.420 | **2.2E-11** | 0.235 | **1.5E-2** | 0.382 | **9.9E-6** | 0.370 | **5.7E-9** | 0.178 | 6.7E-2 | 0.317 | **2.9E-4** |
| PrecentralGMVol vs CSA-GM | 0.119 | 6.9E-2 | 0.080 | 4.1E-1 | 0.176 | **4.8E-2** | 0.102 | 1.2E-1 | 0.092 | 3.5E-1 | 0.141 | 1.2E-1 |
| PostcentralGMVol vs CSA-SC | 0.240 | **2.3E-4** | 0.146 | 1.3E-1 | 0.184 | **4.1E-2** | 0.218 | **8.1E-4** | 0.120 | 2.2E-1 | 0.160 | 7.5E-2 |
| PostcentralGMVol vs CSA-WM | 0.389 | **8.0E-10** | 0.243 | **1.2E-2** | 0.356 | **4.2E-5** | 0.345 | **6.6E-8** | 0.181 | 6.1E-2 | 0.311 | **3.9E-4** |
| PostcentralGMVol vs CSA-GM | 0.034 | 6.1E-1 | 0.008 | 9.3E-1 | 0.059 | 5.1E-1 | 0.064 | 3.3E-1 | 0.050 | 6.1E-1 | 0.088 | 3.2E-1 |
| Cortical Thickness vs CSA-SC | 0.067 | 3.1E-1 | 0.077 | 4.3E-1 | 0.048 | 6.0E-1 | 0.044 | 5.1E-1 | 0.046 | 6.3E-1 | 0.031 | 7.4E-1 |
| Cortical Thickness vs CSA-WM | 0.154 | **1.9E-2** | 0.184 | 5.7E-2 | 0.132 | 1.4E-1 | 0.092 | 1.6E-1 | 0.096 | 3.3E-1 | 0.088 | 3.3E-1 |
| Cortical Thickness vs CSA-GM | -0.069 | 2.9E-1 | -0.051 | 6.0E-1 | -0.086 | 3.4E-1 | 0.029 | 6.6E-1 | 0.090 | 3.5E-1 | -0.027 | 7.6E-1 |
| PrecentralG Thickness vs CSA-SC | 0.211 | **1.3E-3** | 0.180 | 6.3E-2 | 0.201 | **2.5E-2** | 0.144 | **2.8E-2** | 0.125 | 2.0E-1 | 0.117 | 2.0E-1 |
| PrecentralG Thickness vs CSA-WM | 0.252 | **9.9E-5** | 0.249 | **9.7E-3** | 0.209 | **1.9E-2** | 0.146 | **2.5E-2** | 0.142 | 1.5E-1 | 0.087 | 3.3E-1 |
| PrecentralG Thickness vs CSA-GM | 0.113 | 8.5E-2 | 0.083 | 4.0E-1 | 0.149 | 9.6E-2 | 0.081 | 2.2E-1 | 0.122 | 2.1E-1 | 0.044 | 6.2E-1 |
| PostcentralG Thickness vs CSA-SC | 0.035 | 6.0E-1 | 0.145 | 1.3E-1 | -0.013 | 8.8E-1 | 0.014 | 8.3E-1 | 0.116 | 2.3E-1 | -0.027 | 7.6E-1 |
| PostcentralG Thickness vs CSA-WM | 0.152 | **2.1E-2** | 0.245 | **1.1E-2** | 0.171 | 5.6E-2 | 0.084 | 2.0E-1 | 0.165 | 9.0E-2 | 0.108 | 2.3E-1 |
| PostcentralG Thickness vs CSA-GM | -0.069 | 2.9E-1 | 0.018 | 8.5E-1 | -0.160 | 7.4E-2 | 0.062 | 3.4E-1 | 0.152 | 1.2E-1 | -0.030 | 7.4E-1 |
| BrainVol vs Age | -0.079 | 2.3E-1 | -0.165 | 8.4E-2 | -0.150 | 9.0E-2 | | | | | | |
| BrainVol vs Height | 0.612 | **1.2E-25** | 0.274 | **3.8E-3** | 0.408 | **2.1E-6** | | | | | | |
| BrainVol vs Weight | 0.422 | **1.1E-11** | 0.057 | 5.6E-1 | 0.116 | 1.9E-1 | | | | | | |
| BrainGMVol vs Age | -0.188 | **3.7E-3** | -0.309 | **1.0E-3** | -0.264 | **2.6E-3** | | | | | | |
| BrainGMVol vs Height | 0.622 | **1.1E-26** | 0.321 | **6.3E-4** | 0.446 | **1.7E-7** | | | | | | |
| BrainGMVol vs Weight | 0.394 | **2.9E-10** | 0.101 | 2.9E-1 | 0.070 | 4.3E-1 | | | | | | |
| CorticalGMVol vs Age | -0.213 | **9.7E-4** | -0.357 | **1.3E-4** | -0.258 | **3.3E-3** | | | | | | |
| CorticalGMVol vs Height | 0.583 | **6.9E-23** | 0.252 | **7.9E-3** | 0.448 | **1.4E-7** | | | | | | |
| CorticalGMVol vs Weight | 0.351 | **2.6E-8** | 0.041 | 6.7E-1 | 0.069 | 4.4E-1 | | | | | | |

| | | | | | | |
|---|---|---|---|---|---|---|
| CorticalWMVol vs Age | 0.034 | 6.0E-1 | 0.005 | 9.6E-1 | -0.019 | 8.3E-1 |
| CorticalWMVol vs Height | 0.523 | **6.1E-18** | 0.157 | 1.0E-1 | 0.312 | **3.7E-4** |
| CorticalWMVol vs Weight | 0.395 | **2.6E-10** | 0.023 | 8.1E-1 | 0.139 | 1.2E-1 |
| SubCortGMVol vs Age | 0.006 | 9.3E-1 | -0.003 | 9.8E-1 | -0.079 | 3.8E-1 |
| SubCortGMVol vs Height | 0.522 | **7.1E-18** | 0.107 | 2.6E-1 | 0.287 | **1.1E-3** |
| SubCortGMVol vs Weight | 0.426 | **6.8E-12** | 0.169 | 7.8E-2 | 0.078 | 3.8E-1 |
| ThalamusVol vs Age | -0.085 | 1.9E-1 | -0.012 | 9.0E-1 | -0.233 | **8.3E-3** |
| ThalamusVol vs Height | 0.387 | **7.6E-10** | 0.031 | 7.5E-1 | 0.171 | 5.6E-2 |
| ThalamusVol vs Weight | 0.302 | **2.1E-6** | 0.041 | 6.7E-1 | 0.009 | 9.2E-1 |
| CerebellumVol vs Age | -0.120 | 6.5E-2 | -0.097 | 3.1E-1 | -0.260 | **3.0E-3** |
| CerebellumVol vs Height | 0.546 | **1.0E-19** | 0.411 | **8.3E-6** | 0.218 | **1.4E-2** |
| CerebellumVol vs Weight | 0.354 | **2.0E-8** | 0.095 | 3.2E-1 | 0.003 | 9.8E-1 |
| BrainStemVol vs Age | 0.094 | 1.5E-1 | 0.076 | 4.3E-1 | 0.053 | 5.5E-1 |
| BrainStemVol vs Height | 0.531 | **1.6E-18** | 0.310 | **9.8E-4** | 0.258 | **3.6E-3** |
| BrainStemVol vs Weight | 0.431 | **3.6E-12** | 0.119 | 2.1E-1 | 0.191 | **3.1E-2** |
| PrecentralGMVol vs Age | -0.205 | **1.4E-3** | -0.326 | **5.2E-4** | -0.232 | **8.3E-3** |
| PrecentralGMVol vs Height | 0.495 | **4.9E-16** | 0.092 | 3.4E-1 | 0.418 | **1.0E-6** |
| PrecentralGMVol vs Weight | 0.289 | **5.7E-6** | -0.114 | 2.4E-1 | 0.102 | 2.5E-1 |
| PostcentralGMVol vs Age | -0.118 | 6.8E-2 | -0.174 | 7.0E-2 | -0.145 | 1.0E-1 |
| PostcentralGMVol vs Height | 0.434 | **2.8E-12** | 0.121 | 2.1E-1 | 0.369 | **2.0E-5** |
| PostcentralGMVol vs Weight | 0.231 | **3.1E-4** | -0.089 | 3.6E-1 | 0.080 | 3.7E-1 |
| Cortical Thickness vs Age | -0.274 | **1.9E-5** | -0.278 | **3.3E-3** | -0.277 | **1.5E-3** |
| Cortical Thickness vs Height | 0.089 | 1.7E-1 | 0.115 | 2.3E-1 | 0.092 | 3.1E-1 |
| Cortical Thickness vs Weight | -0.020 | 7.6E-1 | 0.010 | 9.1E-1 | -0.083 | 3.5E-1 |
| PrecentralG Thickness vs Age | -0.169 | **8.9E-3** | -0.245 | **9.8E-3** | -0.119 | 1.8E-1 |
| PrecentralG Thickness vs Height | 0.154 | **1.8E-2** | 0.082 | 4.0E-1 | 0.119 | 1.8E-1 |
| PrecentralG Thickness vs Weight | 0.074 | 2.6E-1 | -0.012 | 9.0E-1 | 0.019 | 8.3E-1 |
| PostcentralG Thickness vs Age | -0.149 | **2.2E-2** | -0.097 | 3.1E-1 | -0.187 | **3.4E-2** |
| PostcentralG Thickness vs Height | 0.041 | 5.3E-1 | 0.146 | 1.3E-1 | 0.161 | 7.1E-2 |
| PostcentralG Thickness vs Weight | -0.094 | 1.5E-1 | -0.041 | 6.7E-1 | -0.032 | 7.1E-1 |
