## Supplementary Slides for "Body size interacts with the structure of the central nervous system: A multi-center in vivo neuroimaging study"

### Figure 1 – stable correlation coefficients

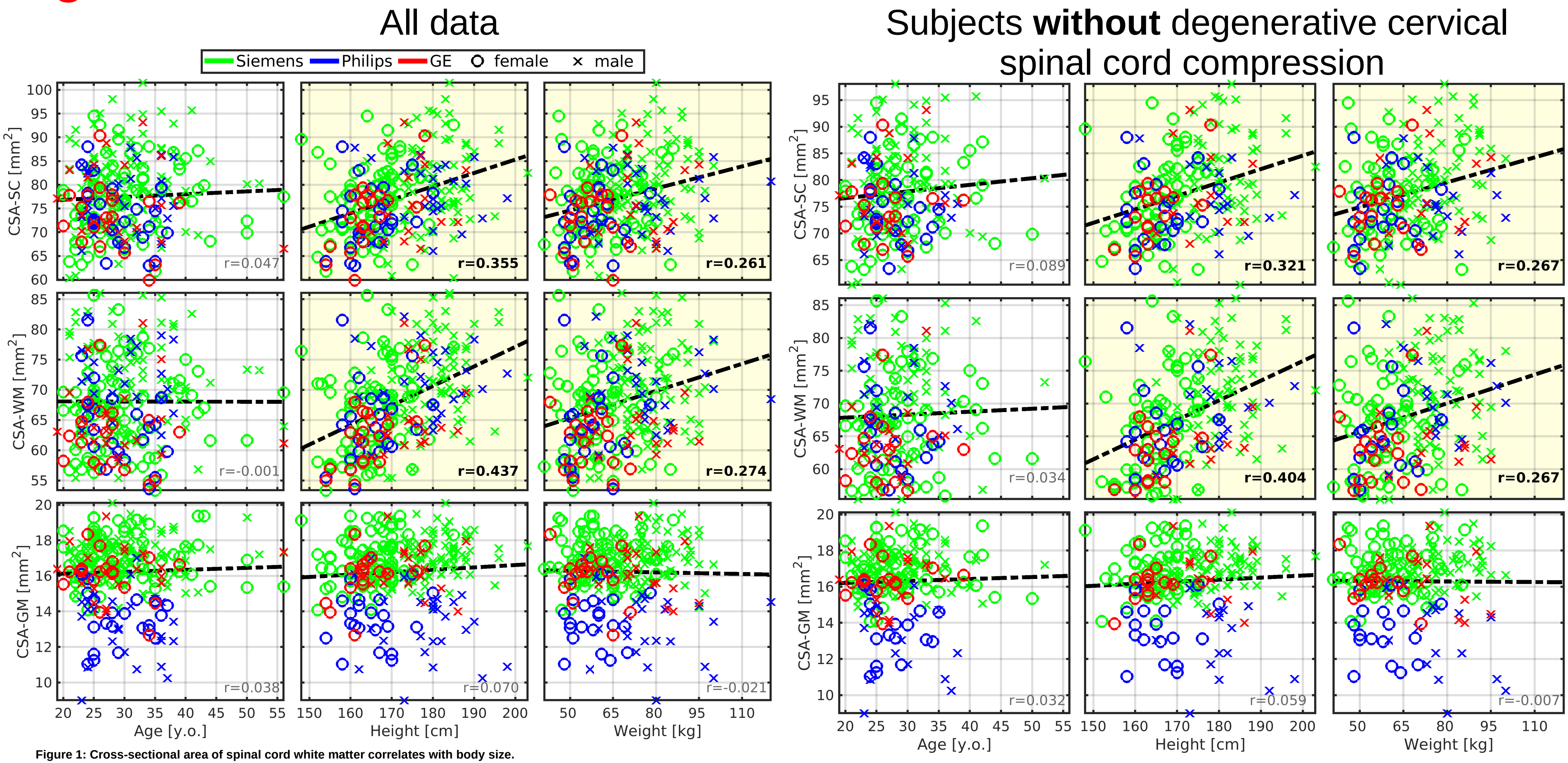


**Figure 1: Cross-sectional area of spinal cord white matter correlates with body size.**
Abbreviations: CSA - cross-sectional area; SC - spinal cord; WM - white matter; GM - gray matter; r - Pearson correlation coefficient. All spinal cord measurements were averaged from cervical C3-4 levels. Regression lines (i.e., the dashed black lines) were estimated from all available data points. Plots with statistically significant correlation (pFWE<0.05) are highlighted with yellow background, and corresponding r values are highlighted with black bold font.

### Figure 1 – stable correlation coefficients

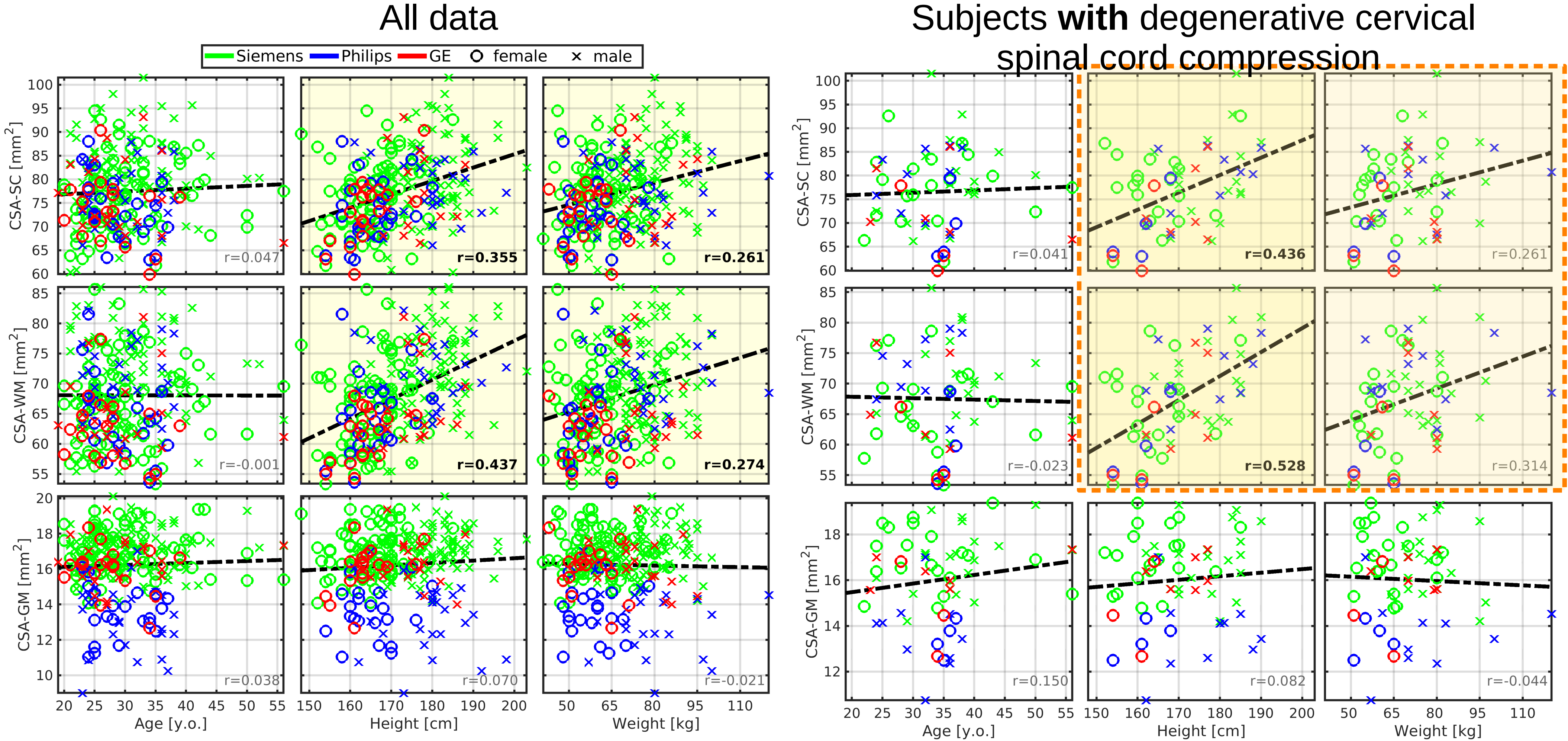


**Figure 1: Cross-sectional area of spinal cord white matter correlates with body size.**
Abbreviations: CSA - cross-sectional area; SC - spinal cord; WM - white matter; GM - gray matter; r - Pearson correlation coefficient. All spinal cord measurements were averaged from cervical C3-4 levels. Regression lines (i.e., the dashed black lines) were estimated from all available data points. Plots with statistically significant correlation (pFWE<0.05) are highlighted with yellow background, and corresponding r values are highlighted with black bold font.

### Figure 3 – stable correlation coefficients

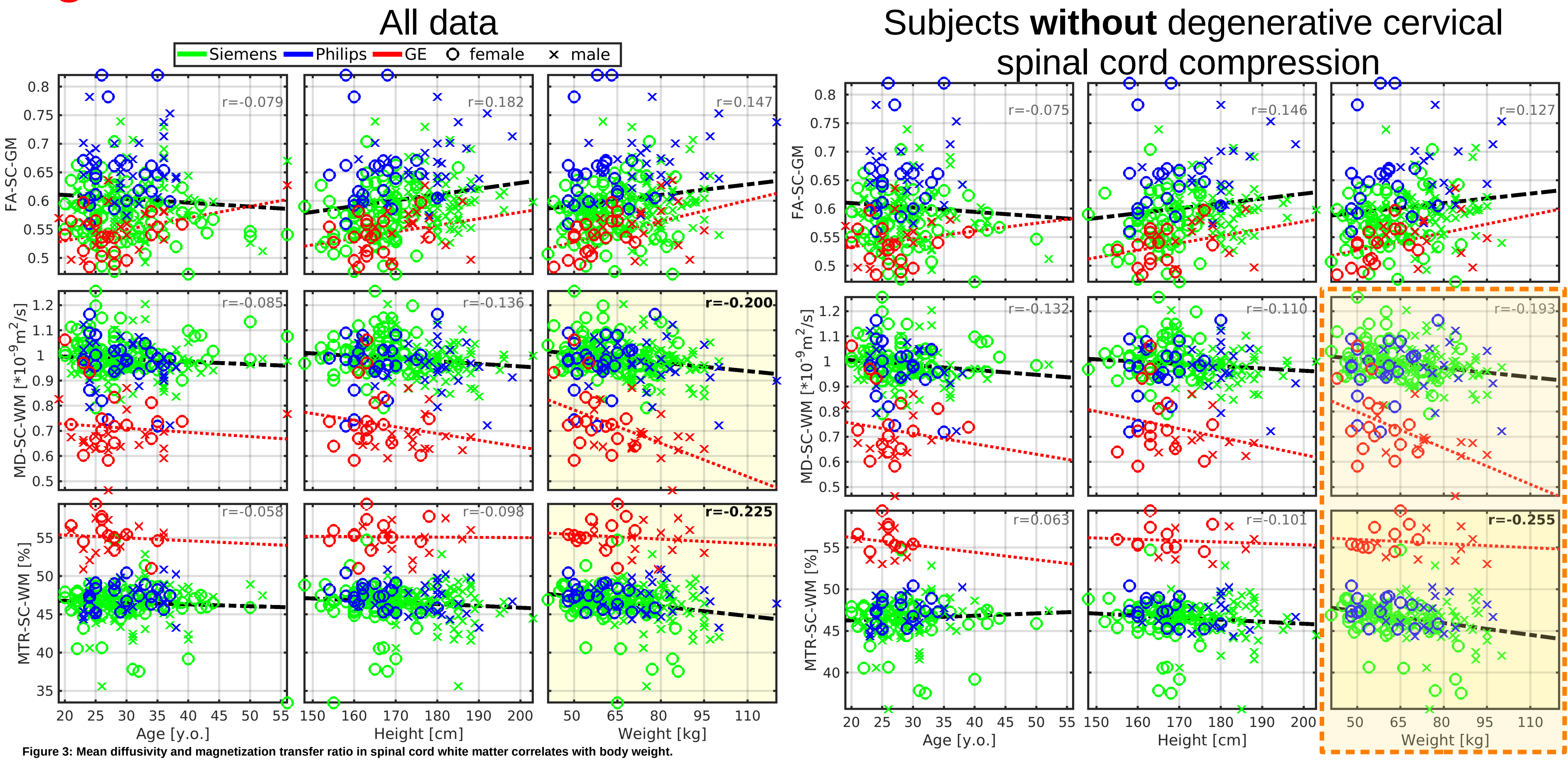


**Figure 3: Mean diffusivity and magnetization transfer ratio in spinal cord white matter correlates with body weight.**
Abbreviations: WM- white matter; MD - mean diffusivity; RD - radial diffusivity; MTR - magnetization transfer ratio; r - Pearson correlation coefficient. All spinal cord measurements were averaged from cervical C2-5 levels. Black dashed regression lines were estimated from the Siemens and Philips scanners' data points. Red dotted regression lines were estimated from the GE scanner's data points. Plots with statistically significant correlation (pFWE<0.05) are highlighted with yellow background, and corresponding r values are highlighted with black bold font.

### Figure 3 – mostly stable correlation coefficients

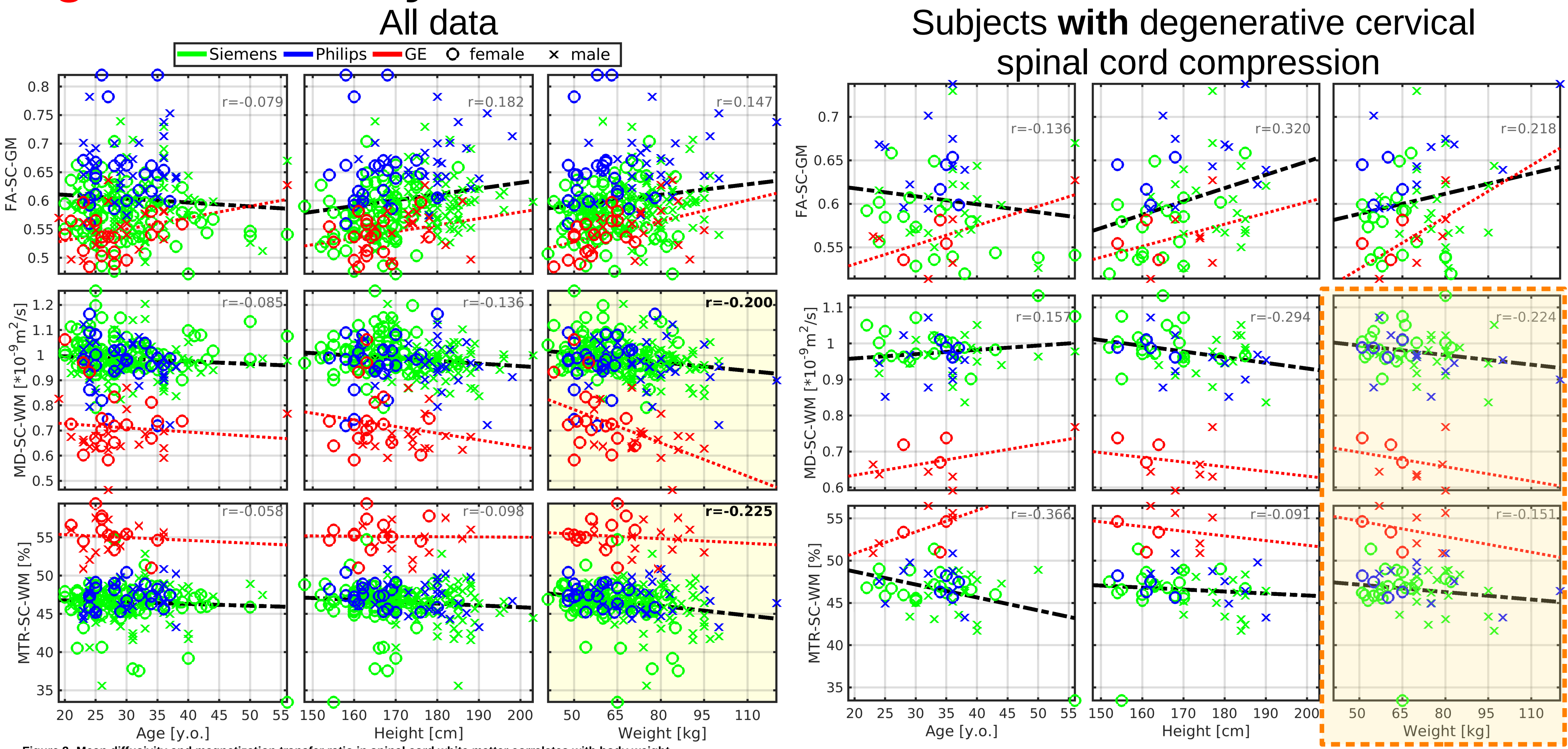


**Figure 3: Mean diffusivity and magnetization transfer ratio in spinal cord white matter correlates with body weight.**
Abbreviations: WM- white matter; MD - mean diffusivity; RD - radial diffusivity; MTR - magnetization transfer ratio; r - Pearson correlation coefficient. All spinal cord measurements were averaged from cervical C2-5 levels. Black dashed regression lines were estimated from the Siemens and Philips scanners' data points. Red dotted regression lines were estimated from the GE scanner's data points. Plots with statistically significant correlation (pFWE<0.05) are highlighted with yellow background, and corresponding r values are highlighted with black bold font.

### **Figure 5b** – stable correlation coefficients

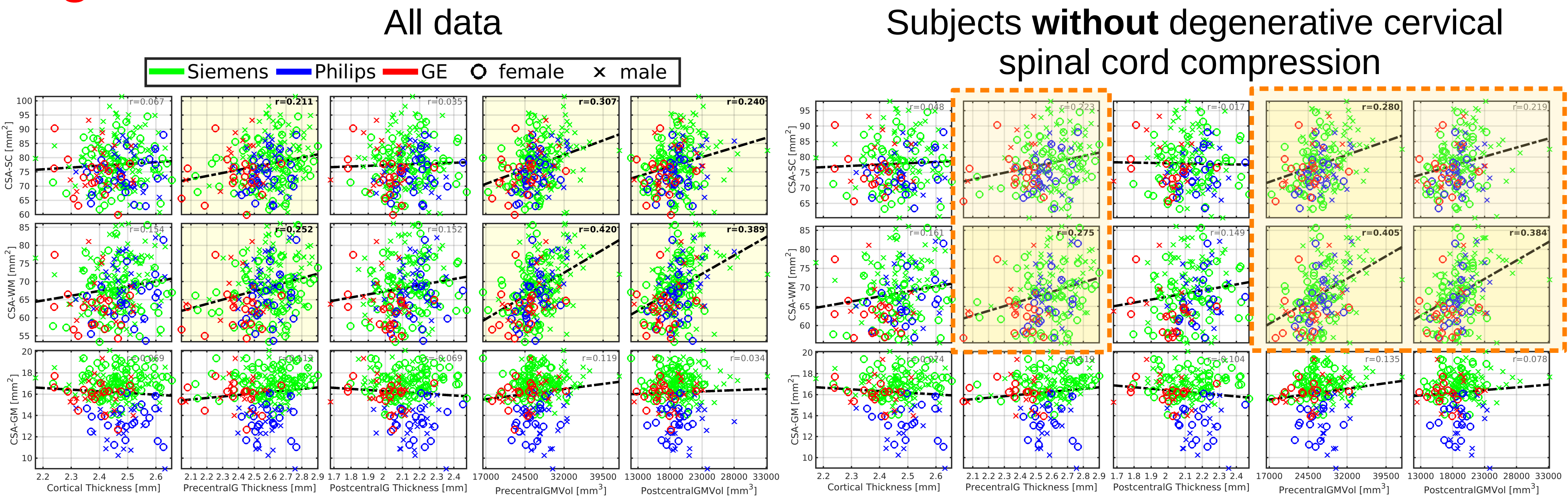


**Figure 5: Cortical morphology correlates with body size, age, and cross-sectional area of the spinal cord white matter.**
Abbreviations: CSA - cross-sectional area; SC - spinal cord; WM - white matter; GM - gray matter; PrecentralG - precentral gyrus; PostcentralG - postcentral gyrus; Vol - volume; r - Pearson correlation coefficient. Regression lines (i.e., the dashed black lines) were estimated from all available data points. Plots with statistically significant correlation (pFWE<0.05) are highlighted with yellow background, and corresponding r values are highlighted with black bold font. Graphs demonstrate correlation with CSA measured in the SC region as averages from cervical C3-4 levels.

### Figure 5b – stable correlation coefficients

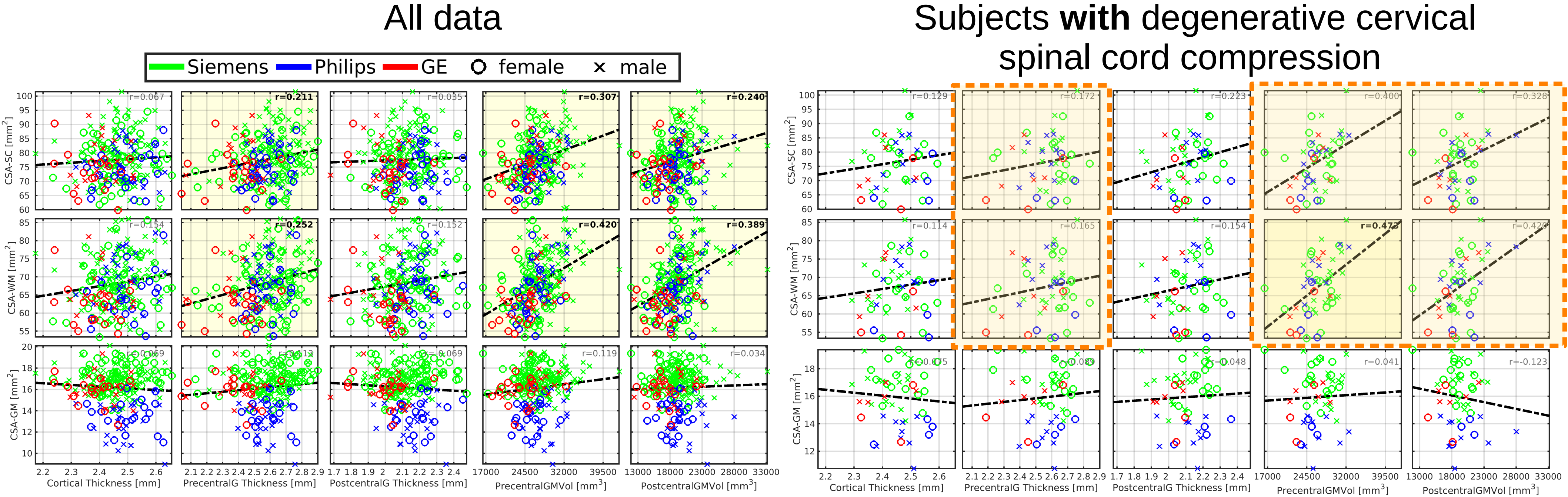


**Figure 5: Cortical morphology correlates with body size, age, and cross-sectional area of the spinal cord white matter.**
Abbreviations: CSA - cross-sectional area; SC - spinal cord; WM - white matter; GM - gray matter; PrecentralG - precentral gyrus; PostcentralG - postcentral gyrus; Vol - volume; r - Pearson correlation coefficient. Regression lines (i.e., the dashed black lines) were estimated from all available data points. Plots with statistically significant correlation (pFWE<0.05) are highlighted with yellow background, and corresponding r values are highlighted with black bold font. Graphs demonstrate correlation with CSA measured in the SC region as averages from cervical C3-4 levels.

### Figure 6 – stable correlation coefficients

#### All data

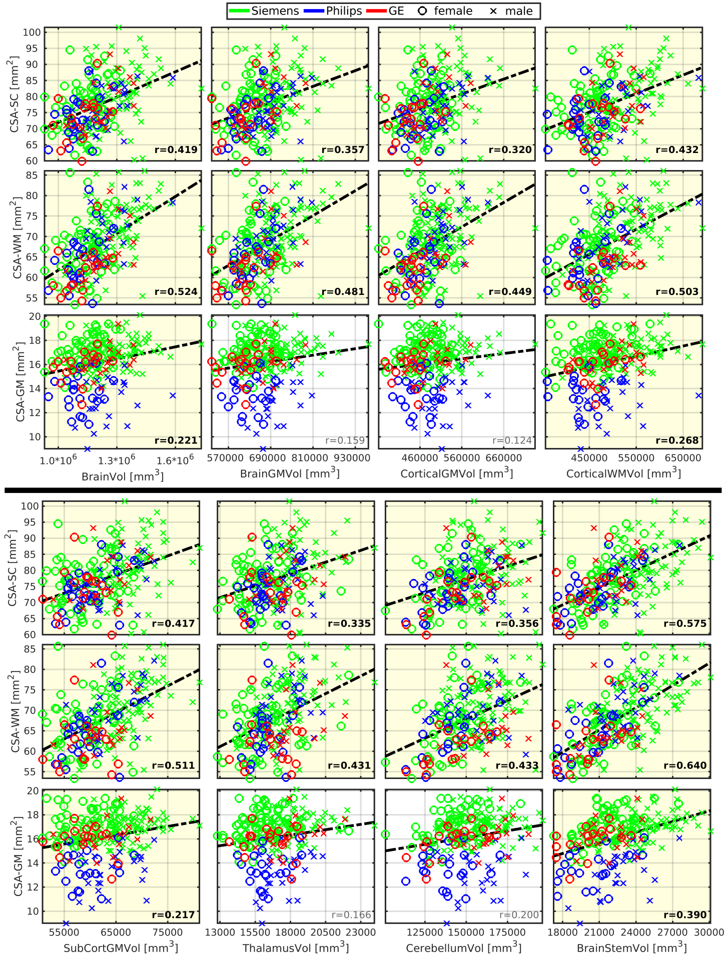


#### Subjects **without** degenerative cervical spinal cord compression

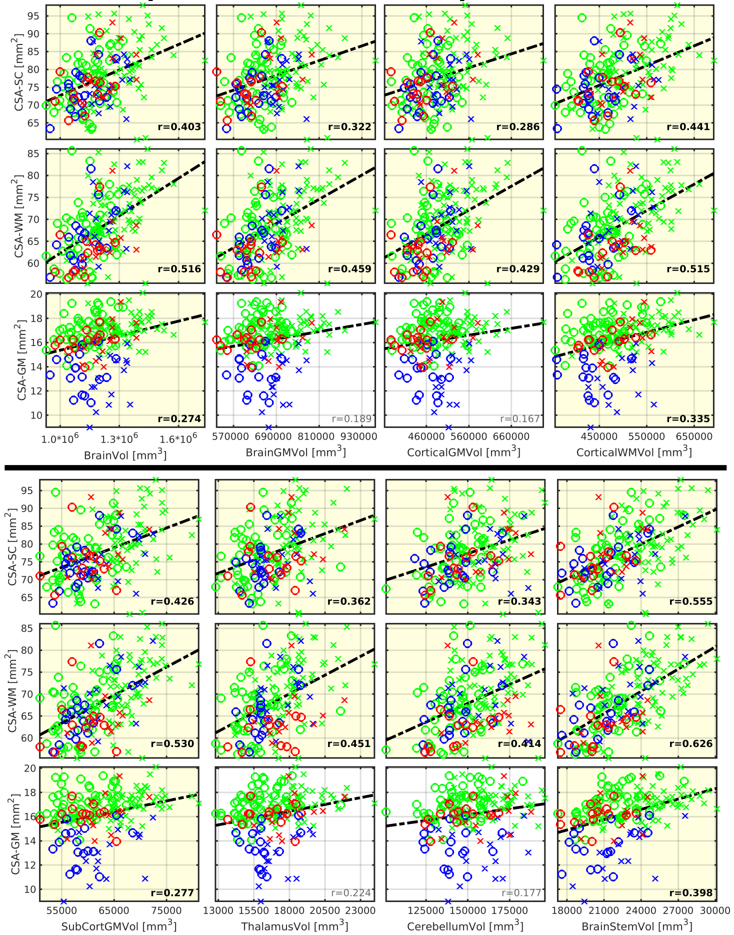


**Figure 6: Brain morphology correlates with spinal cord morphology.** Abbreviations: CSA - cross-sectional area; SC - spinal cord; WM - white matter; GM - gray matter; Vol - volume; SubCort - subcortical; r - Pearson correlation coefficient. All SC measurements were averaged from cervical C3-4 levels. Regression lines (i.e., the dashed black lines) were estimated from all available data points. Plots with statistically significant correlation (pFWE<0.05) are highlighted with yellow background, and corresponding r values are highlighted with black bold font.

### Figure 6 – stable correlation coefficients

#### All data

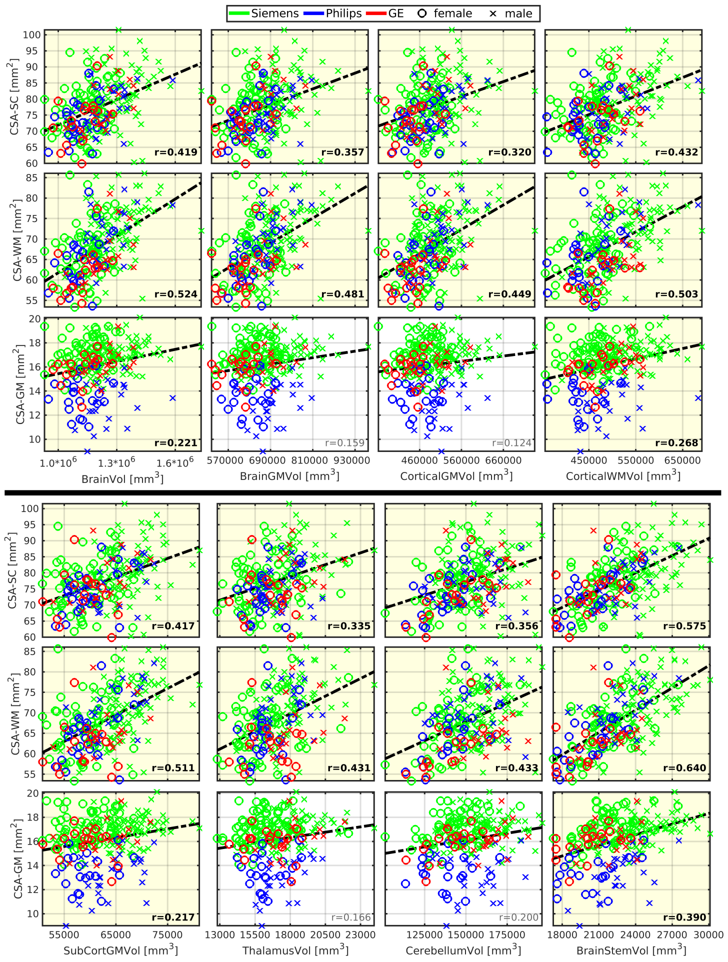


**Figure 6: Brain morphology correlates with spinal cord morphology.** Abbreviations: CSA - cross-sectional area; SC - spinal cord; WM - white matter; GM - gray matter; Vol - volume; SubCort - subcortical; r - Pearson correlation coefficient. All SC measurements were averaged from cervical C3-4 levels. Regression lines (i.e., the dashed black lines) were estimated from all available data points. Plots with statistically significant correlation (pFWE<0.05) are highlighted with yellow background, and corresponding r values are highlighted with black bold font.

#### Subjects **with** degenerative cervical spinal cord compression

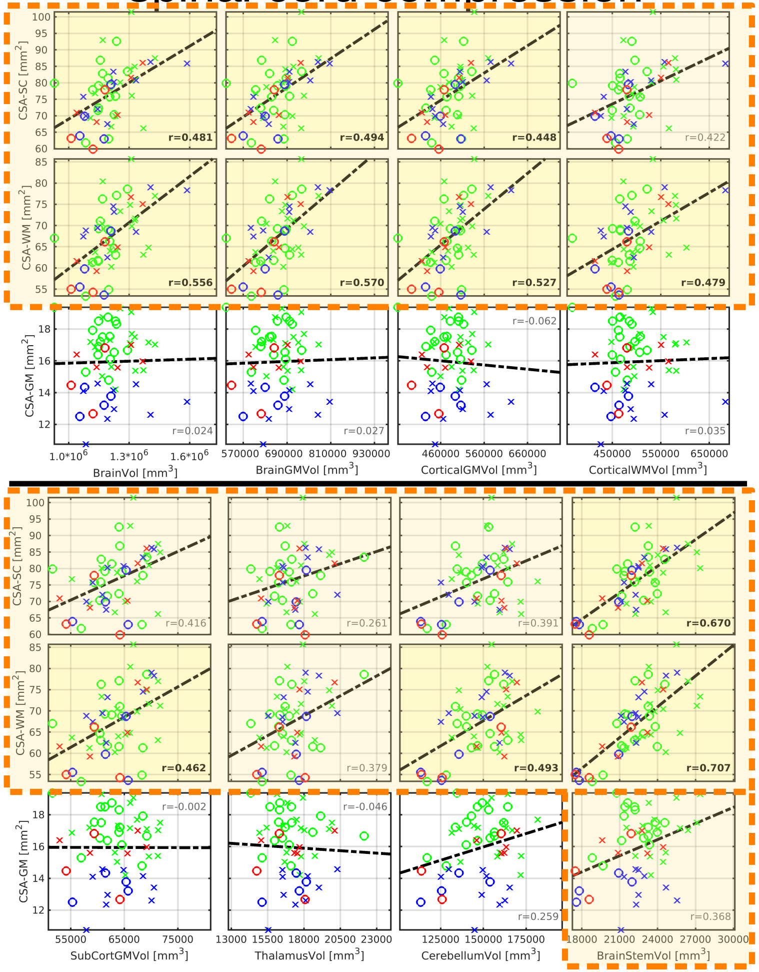
